# The spatio-temporal dynamics of phoneme encoding in aging and aphasia

**DOI:** 10.1101/2024.10.21.619562

**Authors:** Jill Kries, Maaike Vandermosten, Laura Gwilliams

## Abstract

During successful language comprehension, speech sounds (phonemes) are encoded within a series of neural patterns that evolve over time. Here we tested whether these neural dynamics of speech encoding are altered for individuals with a language disorder. We recorded EEG responses from individuals with post-stroke aphasia and healthy age-matched controls (i.e., older adults) during 25 minutes of natural story listening. We estimated the duration of phonetic feature encoding, speed of evolution across neural populations, and the spatial location of encoding over EEG sensors. First, we establish that phonetic features are robustly encoded in EEG responses of healthy older adults. Second, when comparing individuals with aphasia to healthy controls, we find significantly decreased phonetic encoding in the aphasic group after shared initial processing pattern (0.08-0.25s after phoneme onset). Phonetic features were less strongly encoded over left-lateralized electrodes in the aphasia group compared to controls, with no difference in speed of neural pattern evolution. Finally, we observed that healthy controls, but not individuals with aphasia, encode phonetic features longer when uncertainty about word identity is high, indicating that this mechanism - encoding phonetic information until word identity is resolved - is crucial for successful comprehension. Together, our results suggest that aphasia may entail failure to maintain lower-order information long enough to recognize lexical items.

**Significance statement:** This study reveals robust decoding of speech sound properties, so-called phonetic features, from EEG recordings in older adults, as well as decreased phonetic processing in individuals with a language disorder (aphasia) compared to healthy controls. This was most prominent over left-hemispheric electrodes. Additionally, we observed that healthy controls, but not individuals with aphasia, encode phonetic features longer when uncertainty about word identity is high, indicating that this mechanism - encoding phonetic information until word identity is resolved - is crucial for successful language processing. These insights deepen our understanding of disrupted mechanisms in a language disorder, and show how the integration between language processing levels works in the healthy aging, neurotypical brain.

## 1 Introduction

Comprehending natural speech entails rapidly converting continuous acoustic input into discrete units, such as phonemes and words (Hickok and Poeppel, 2007). A study by Gwilliams et al. (2022) demonstrated that the spectrotemporal properties of phonemes, known as phonetic features, are dynamically encoded in the brain. This coding scheme facilitates integration with higher-level language representations, such as syntactic or semantic features, through hierarchical dynamic coding (HDC) (Gwilliams et al., 2024). Individuals with language disorders such as aphasia can have deficits in phoneme identification (Kries et al., 2023) or phonological processing (Blumstein, 1998), among other difficulties. With the HDC framework, we now have a unique opportunity to directly investigate the neural dynamics of speech representations in individuals with aphasia (IWA). This paper aims to address this research gap by investigating the effects of aphasia on phonetic feature encoding.

When healthy young adults listen to speech, phonetic features are encoded in auditory cortex, including the transverse temporal gyrus, superior temporal gyrus, and superior temporal sulcus (Mesgarani et al., 2007, 2014; Wilson et al., 2018; Levy and Wilson, 2019; Yi et al., 2019). Based on magnetoencephalography (MEG) data during natural story listening, Gwilliams et al. (2022) showed that the spectro-temporal properties of the phonetic features are encoded for around 0.3 s, outlasting the duration of the phoneme input, and that the encoded information is passed to a new neural ensemble every 0.08 s, which is proportional to the average phoneme duration in the stimulus (Gwilliams et al., 2022, 2024). The authors conclude that phonetic features are encoded in a dynamic neural pattern, whereby different neural ensembles are recruited across time – referred to as HDC (Gwilliams et al., 2024). This means that rather than a one-to-one mapping between neural ensemble and feature encoding, different ensembles are activated in sequence. This allows a running history of the past three phonemes to be encoded in parallel, while also keeping track of the relative order that they were said.

Aphasia is often considered primarily a phonological, semantic or syntactic language disorder, due to commonly observed symptoms like word finding difficulties or incomplete sentences (Harnish, 2018; Zhang and Hinzen, 2022). However, recent work suggests that the processing issues associated with aphasia can also be related to more low-level speech processing, i.e., neural encoding of amplitude fluctuations of speech and phoneme identification (Kries et al., 2023, 2024). This implies that some language difficulties experienced in aphasia may arise from an impoverished representation of speech sounds. In fact, Kries et al. (2023) found that IWA’s performance at discriminating between different rates of change in amplitude predicted phonological processing performance, suggesting that low-level processing impairments may propagate onto higher-level processes. Building upon this prior work, we aim to assess whether difficulties in phoneme processing in aphasia may stem from challenges in dynamic coding of phonetic features.

Gwilliams et al. (2022) used MEG recordings to investigate phonetic encoding during a two-hour story-listening task in younger adults. By contrast, we are using an available electroencephalography (EEG) dataset of IWA and healthy age-matched controls listening to natural speech for 25 minutes (fig.1B) (Kries et al., 2024). EEG is more feasible than MEG for post-stroke or older adults, because it is more accessible in clinical settings, and allows for more flexible body position during the recording. Moreover, shorter neural recordings are desirable to maintain optimal attention levels. Therefore, our first research question examines whether the robust phonetic decoding observed by Gwilliams et al. (2022) in younger adults can be replicated with (1) EEG data, (2) a shorter recording duration and (3) in older adults.

**Figure 1:**
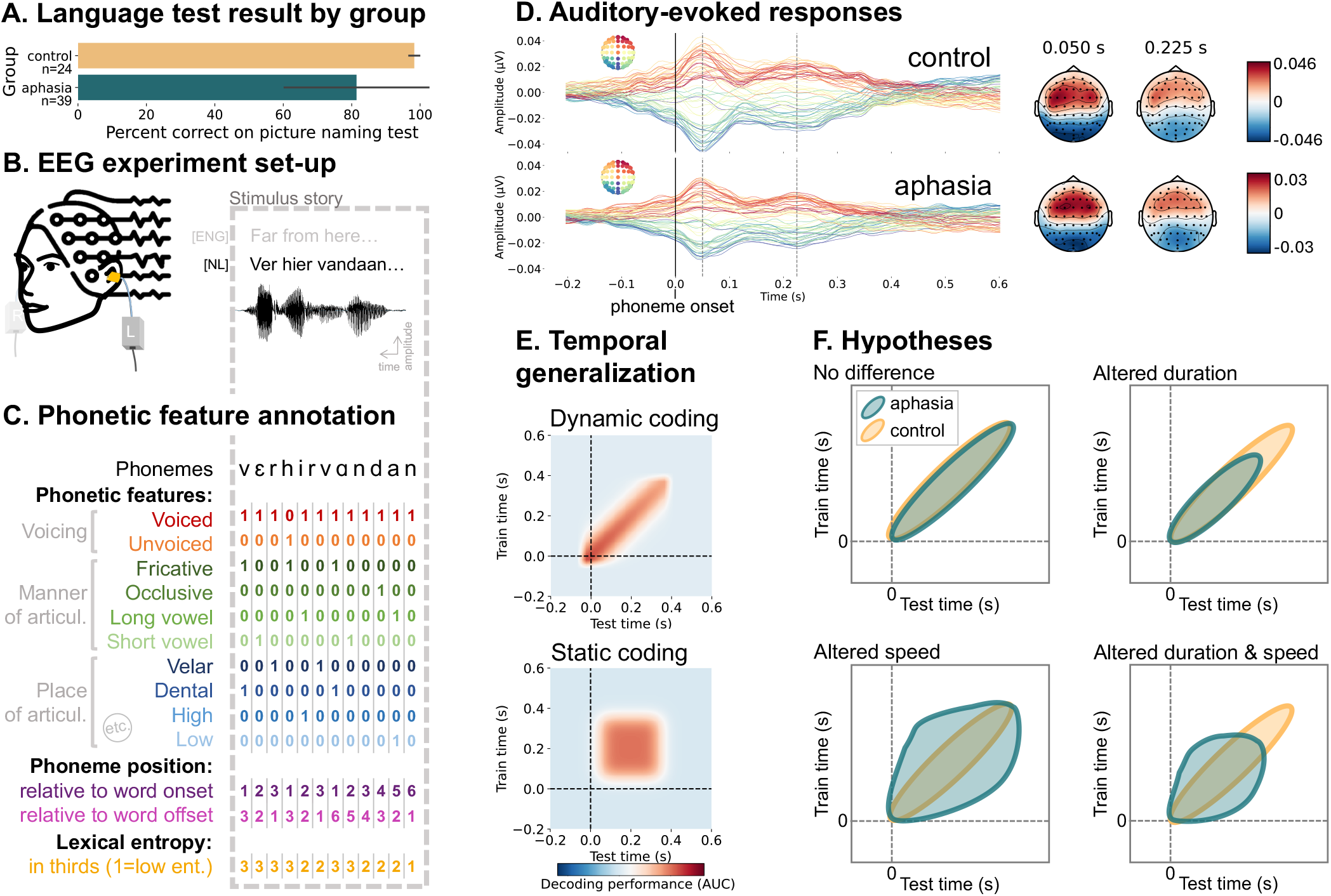
Experimental set-up, schematic of analytic approach, auditory-evoked responses by group and hypothesis illustration. **(A)** Mean and standard deviation for the picture naming test scores (in percentage) by group. **(B)** The experimental protocol consisted of participants listening to a 25 minute-long narrative in Flemish while 64-sensor-EEG data was being recorded. The audio was presented bilaterally. The first 3 words and their acoustic waveform are depicted, with an English translation. **(C)** The narrative was annotated for 18 phonetic features (not all illustrated) with across-feature time alignment. Moreover, the phonemes were subset by position within word and by lexical entropy (entropy values were categorized into thirds). **(D)** The EEG data was epoched around the phoneme onsets. This plot displays the timecourses of the auditory evoked response for all 64 EEG sensors on the left and the topographies at 0.05s and 0.225s after phoneme onset on the right, for the control group and aphasia group separately. **(E)** Temporal generalization: Training a decoder at each time slice and testing each of these decoders across all possible time slices allows to investigate the encoding duration (matrix diagonal) and generalization (width of significant cluster). The top plot shows a dynamic encoding pattern wherein the encoded information evolves across temporal decoders, i.e., the way that information is encoded changes over time. The bottom plot shows a static encoding pattern wherein the encoded information is maintained. **(F)** We hypothesized that the aphasia group would show a different encoding pattern than the healthy control group. The top right graph illustrates what the temporal generalization (TG) pattern would look like if individuals with aphasia had a shorter encoding duration than controls. The bottom left graph shows what we would expect to see if individuals with aphasia had a slower speed of evolution of encoding than controls, i.e., longer generalization. The bottom right graph displays what we would expect to observe if both duration and generalization of encoding would be altered in aphasia.

The second research question examines the dynamics of phonetic encoding in IWA compared to age-matched healthy controls. Specifically, we wanted to know whether IWA would show a change in phonetic encoding duration, a change in speed of neural pattern evolution, or a mix of both (fig.1F). We hypothesized slower speed of neural pattern evolution in aphasia, because that would result in representational overlap between neighbouring phonemes and might explain symptoms observed in aphasia, such as phonemic paraphasias. Given the structural and functional neural changes following stroke-induced aphasia (Wilson et al., 2023), we also test for differences in topography of neural activity during phonetic encoding between IWA and healthy controls.

## 2 Methods and methods

### 2.1 Participants

39 IWA in the chronic phase after stroke (≥ 6 months after onset; age in years: mean(standard deviation)=69.5(12.4)) and 24 demographically-matched healthy control participants (age in years: mean(standard deviation)=71.5(7)) listened to a narrative for 25 minutes (in their native language, Dutch) while EEG data was recorded (fig.1B). IWA were recruited at the University hospital in Leuven and via advertisement, see supplementary section S.11 for more information. All aphasia participants in the used dataset had clinically diagnosed post-stroke aphasia in the acute phase after stroke (0-14 days post-stroke). We only included IWA that had no formal diagnosis of a psychiatric or neurodegenerative disorder and that had a left-hemispheric or bilateral lesion. All aphasia participants were tested in the chronic phase after stroke (time since stroke onset in months (median(range)): 16.1(6-126.1), after spontaneous recovery and therapy. 37/39 participants (95%) still had residual language impairments at the moment of data collection, as determined by clinical evaluation, and evidenced by continued attendance to speech therapy sessions.

The aphasia sample was checked for language impairments at the moment of data collection using two standardized diagnostic aphasia tests, i.e., the diagnostic test ScreeLing (Visch-Brink et al., 2010) and the Dutch picture-naming test (NBT; Van Ewijk et al. (2020)), using the same procedure as reported in Kries et al. (2024) (fig.1A; supplementary table S.2). These tests reflect aphasia severity (also see supplementary section S.4). On both tests, the aphasia group had significantly lower results than the control group (ScreeLing cut-off threshold: 68/72 points; NBT cut-off threshold: 255/276 points; ScreeLing (mean(standard deviation)): aphasia=61.5(9.4), control=69.9(2.5), p<.001; NBT (mean(standard deviation)): aphasia=224.7(58.1), control=271.3(4.1), p<.001; supplementary table S.1). Moreover, all participants completed further behavioral tasks assessing: hearing (pure tone audiogram), rise time discrimination, phoneme identification4, phonological and semantic word fluency, cognition (total score of attention, executive function and memory tasks (Huygelier et al., 2019)), self-reported alertness and fatigue, story comprehension question accuracy (Kries et al., 2024). More details on these tasks and the group comparisons are available in the supplementary material of this paper (supplementary section S.4). 82% of the aphasia group showed deficits at either one of the 2 phonological tasks that we administered (i.e., phoneme identification task and phonology subtest of the ScreeLing; supplementary section S.4).

Throughout the paper, we are referring to healthy control participants and older adults interchangeably given that these participants are on average 71 years old, with a range of 58 to 87 years of age. All participants were Dutch native speakers from Flanders, Belgium. Age, sex, education, handedness and multilinguality did not differ between groups (supplementary table S.1; Kries et al. (2024)). Demographic information was acquired via a self-reported questionnaire. Handedness was assessed via the Edinburgh Handedness Inventory (Oldfield, 1971). Details about the stroke in IWA, i.e., time since stroke onset, stroke type, occluded blood vessel, lesion location and speech-language therapy, can be found in supplementary table S.2. To visualize the damaged brain tissue of IWA, a lesion overlap image was created (supplementary fig.S.1). Informed consent was obtained from all participants for the recruitment via screening and for the data collection in the chronic phase. The study received ethical approval by the medical ethical committee of KU Leuven and UZ Leuven (S60007) and is in accordance with the declaration of Helsinki.

### 2.2 Experimental design

The EEG measurements were conducted in a soundproof room with a Faraday cage using a 64-channel EEG system (ActiveTwo, BioSemi, Amsterdam, NL) at 8192 Hz sampling frequency. Participants listened to a 25-minute story, The Wild Swans written by Hans Christian Andersen, narrated by a female Flemish-native speaker. The stimulus story was specifically tailored to the aphasia population by choosing a fairytale as stimulus that consists of simple language. The narrative comprised 13560 phonemes and 3002 words. Stimuli were presented binaurally via shielded insert earphones at 60 dB SPL (A weighted). Participants were asked 5 yes/no and 5 multiple-choice questions about the story content throughout to ensure attentiveness (see supplementary section S.4 for more information).

### 2.3 EEG signal processing

The EEG signal processing was conducted using MATLAB (version 9.1.0.441655 (R2016b)). Eye movement artifact removal was employed using the multichannel Wiener filter (Somers et al., 2018). The EEG signal was referenced to the common average. For high-pass filtering, we applied a least squares filter with filter order of 2000, passband frequency of 0.5 Hz and stopband frequency of 0.45 Hz. For low-pass filtering, we employed a least squares filter with a filter order of 2000, passband frequency of 25 Hz and stopband frequency of 27.5 Hz. Subsequently, EEG data was downsampled to 128 Hz and normalized by subtracting the mean and dividing by the standard deviation. Next, the EEG data was epoched, centered around phoneme onsets, specifically from -0.2s before phoneme onset to 0.6s after onset. No baseline correction was applied during epoching.

### 2.4 Modelled features

The stimulus narrative was annotated for different properties at the phoneme level, namely 21 phonetic features, phoneme position relative to word onset and relative to word offset, as well as lexical entropy (fig.1C). To achieve this, an aligner (Duchateau et al., 2009) was used to create alignment files containing the identity and timing of each phoneme and each word of the stimulus. Then, we annotated phonetic features based on the multi-value feature system reported in (King and Taylor, 2000), which features voicing, manner of articulation, place of articulation, roundness and front-backness. Voicing indicates whether or not the vocal cords vibrate during speech production. **Voicing** has 2 features, voiced (e.g., /b/) and unvoiced (e.g., /p/). **Manner** describes how air flows through the articulators during speech production. We tested 6 manner features: nasal, fricative, occlusive and approximant for consonsants, and short vowel and long vowel for vowels. **Place** refers to the positioning of the articulators (e.g., teeth, tongue, lips) during speech production. We tested 8 place features: dental, coronal, glottal, labial and velar for consonsants, and low, mid, high for vowels. **Roundness** describes whether or not lips are rounded during pronunciation, thus consisting of 2 features, rounded (e.g., /w/) and unrounded (e.g., /f/). **Front-backness** refers to the position of the tongue in the mouth, entailing the features front, central and back. We removed phonetic features that occurred less than 5% of total phonemes in the stimulus, leading to removal of 3 place features, i.e., dental, glottal and low vowel. This led to 18 features that were used for further analysis.

#### 2.4.1 Subset variables

To investigate whether phonemes are differentially encoded depending on their phoneme position within words and their lexical entropy, we subset phonemes based on these properties. **Phoneme position** relative to word onset was coded as 1 for the first phoneme within words, 2 for the second phoneme within words, etc. until the 5th position, and the opposite order for phoneme position relative to word offset. This means that there were less phoneme occurrences per position, as evidenced in table 1. Concretely, this was executed by using the word onsets to chunk the phoneme onsets by word, then loop through the chunks and serialize the phoneme onsets within word. **Lexical entropy** measures the level of competition among words that match the current phoneme input. For example, upon hearing the sounds /pl/, many potential words are activated (n=999), indicating a high degree of competition. As additional phonemes are heard, the number of possible words diminishes (e.g., for /plu/, n=162, and for /plur/, n=19), resulting in a lower degree of competition. This competition level is calculated using the Shannon entropy of the words in the activated cohort (Gwilliams and Davis, 2022). The first phoneme of each word was excluded from this analysis. Entropy was calculated based on the SUBTLEX-NL database (Keuleers et al., 2010) and a custom pronunciation dictionary. Then we computed the 33rd and 66th percentiles to split entropy values into 3 equal parts and used the third with the lowest and the third with the highest values for the decoding analysis. Table 1 shows how many phoneme occurrences were present in the thirds.

**Table 1:** Phoneme occurrences for subset variables.

| level | n | meaning |
| --- | --- | --- |
| phoneme position relative to word onset/offset |  |  |
| 1 | 3002 | first/last phoneme within word |
| 2 | 2948 | second (to last) phoneme within word |
| 3 | 2199 | third (to last) phoneme within word |
| 4 | 1524 | fourth (to last) phoneme within word |
| 5 | 1187 | fifth (to last) phoneme within word |
| lexical entropy |  |  |
| 1 | 3427 | phonemes with low entropy |
| 3 | 3592 | phonemes with high entropy |

### 2.5 Decoding analysis

We decoded each phonetic feature from the 64-sensor EEG signal. All decoding was performed using one- versus-all logistic regression and 5-fold cross-validation, using a temporal generalization (TG) approach. TG entails testing whether a temporal decoder trained on data at time t can accurately decode data at time t from a testing set (fig.1E). Instead of assessing decoding accuracy solely at the specific time point the model was trained on, we evaluated its accuracy across all possible train/test time combinations. TG is represented by a square decoding matrix of training time vs. testing time (fig.1E). We assessed the decoding performance by computing the area under the curve (AUC) of the receiver-operating curve.

To assess for how long phonetic features can be decoded from brain data, we extracted the temporal decoders where train and test time were identical, i.e., the diagonal of the TG matrix, and extracted above-chance time points. To assess for how long information captured at any given temporal decoder is maintained or generalized, we extracted the width of above-chance clusters of the TG full matrix. Additionally, to explore the dynamics of neural representations, we compared the mean duration of above-chance generalization across temporal decoders to the duration of above-chance temporal decoding (i.e., comparing the rows of the matrix to its diagonal). These metrics were examined within each participant and analyzed using second-level statistics across participants.

To investigate which EEG sensors are most important for phonetic decoding, we decoded features from each sensor individually, thus passing the epochs’ timecourse of a given sensor to the decoder. This can tell us how much each sensor contributes across time to phonetic feature decoding. Note on decoding/encoding terminology: we employ decoding analyses, using EEG data as input to predict speech features. Successfully decoding a feature implies its encoding in neural activity. We chose decoding because it (i) replicates the previous study’s approach (Gwilliams et al., 2022) and (ii) uses multivariate EEG data, offering robustness against noise and aggregating signals across channels (Brodbeck et al., 2021).

#### 2.5.1 Comparing performance between trial subsets

To assess decoding performance across theoretically interesting subsets of trials (such as start vs. end of a word or high vs. low entropy), we slightly changed our train/test cross-validation procedure. The classifier was trained using the full training dataset, while the test set was divided according to the different levels of interest (table 1). We independently evaluated the model’s performance on each subset of the test data, yielding a timecourse or generalization matrix for each group of trials being analyzed.

### 2.6 Statistical analysis

To compare TG matrices and the derived diagonals against chance-level, we used permutation-based statistics. This was performed on group-level average decoding performance using a one-sample permutation cluster test implemented using MNE Python (Gramfort et al., 2013). First, a t-value is calculated for every data point of the TG matrix. To determine statistical significance, the data undergoes 10,000 iterations of random permutation, wherein the signs of data points are randomly flipped. For each permutation, clusters of neighboring data points are identified, and the sum of t-values within these clusters is computed. This generates a null distribution of summed t-values from the permuted data. The significance of the observed effect is then assessed by comparing the observed cluster’s t-value sum to this null distribution. If the observed sum is greater than 95% of sums from the null distribution, the effect is considered statistically significant.

To compare aphasia and control groups, we used the permutation cluster test as implemented in MNE Python (Gramfort et al., 2013). This test compares groups by first calculating differences at each data point of the TG matrices. It then shuffles data between groups 10’000 times to create a null distribution. Significant clusters, where differences are consistent, are identified by comparing observed values to those in the null distribution.

To test whether the TG matrix pattern is dynamically evolving (i.e., the diagonal is larger than the width of the decoding pattern), we (1) extracted the diagonal and horizontal (average across y-axis) timecourses per participant, (2) subtracted the diagonal from the horizontal timecourse for each participant, and (3) used a one-sample permutation cluster test to test whether this delta is significantly different from zero.

To analyze topographical differences of phonetic feature decoding between aphasia and control groups, we submitted the decoding performance of each sensor across subjects and groups to a mass-univariate independent samples t-test (Brodbeck, 2020).

## 3 Results

### 3.1 Replicating phonetic encoding dynamics in older adults with EEG data

Gwilliams et al. (2022) found that the time window in which phonetic features were significantly decodable from MEG data was from 0.05 to 0.3s after phoneme onset. Using the same approach, we computed a timecourse of decoding performance, reflecting to what extent the phonetic information is present in the EEG signal, for each of the 18 tested phonetic features. We found that all 18 phonetic features were decodable above chance using one-sample permutation cluster tests across subjects (cluster-forming threshold: p=.05; table 2 and fig.2A). Due to variations in phoneme duration, such as between vowels and consonants, and the differing number of phoneme trials within certain phonetic feature categories, there is variability in decoding performance between features. Therefore, we focus on the decoding performance averaged across all features moving forward. When averaging across phonetic features, we observed above chance decoding from -0.04 to 0.49s (p<.001) relative to phoneme onset for the control group (fig.2B top right plot). Thus, on average, phonetic features are decodable for a span of 0.53s. Given that the average phoneme duration in the story was 0.098s (standard deviation=0.107s), this replicates the prior finding from younger adults (Gwilliams et al., 2022) that neural encoding of phonetic features long surpasses the duration of the phoneme itself. Results hereafter are all based on the average across phonetic features.

**Table 2:** Above-chance decoding of phonetic features in healthy older adults.

| phonetic feature | significant time window (s) | average decoding performance (AUC) | average t-value | p-value |
| --- | --- | --- | --- | --- |
| voiced | -0.052 to 0.452 | 0.515 | 1.414 | <.001 |
| unvoiced | 0.025 to 0.304 | 0.509 | -0.009 | <.001 |
| nasal | -0.013 to 0.165 | 0.508 | 0.785 | <.001 |
| fricative | 0.102 to 0.165 | 0.508 | -1.418 | 0.003 |
|  | 0.188 to 0.258 | 0.508 | 0.497 | <.001 |
| occlusive | 0.056 to 0.258 | 0.518 | 1.662 | <.001 |
| approximant | 0.064 to 0.157 | 0.508 | 2.42 | <.001 |
| short vowel | -0.013 to 0.258 | 0.514 | 1.029 | <.001 |
| long vowel | -0.099 to 0.374 | 0.518 | 1.258 | <.001 |
| coronal | 0.025 to 0.157 | 0.511 | 1.905 | <.001 |
|  | 0.180 to 0.250 | 0.506 | 1.220 | <.001 |
| labial | 0.079 to 0.141 | 0.516 | 0.758 | 0.002 |
|  | 0.157 to 0.250 | 0.510 | -0.077 | 0.002 |
|  | 0.273 to 0.390 | 0.511 | -0.963 | <.001 |
| velar | -0.153 to -0.114 | 0.506 | 1.059 | 0.009 |
| mid | -0.013 to 0.266 | 0.515 | -0.484 | <.001 |
| high | 0.064 to 0.289 | 0.516 | -1.238 | <.001 |
| rounded | 0.048 to 0.359 | 0.511 | -0.196 | <.001 |
| unrounded | -0.044 to 0.6 | 0.512 | 2.255 | <.001 |
| front | 0.071 to 0.483 | 0.506 | 1.406 | <.001 |
| central | -0.013 to 0.165 | 0.517 | 0.686 | <.001 |
| back | 0.203 to 0.258 | 0.504 | 1.307 | 0.003 |

**Figure 2:**
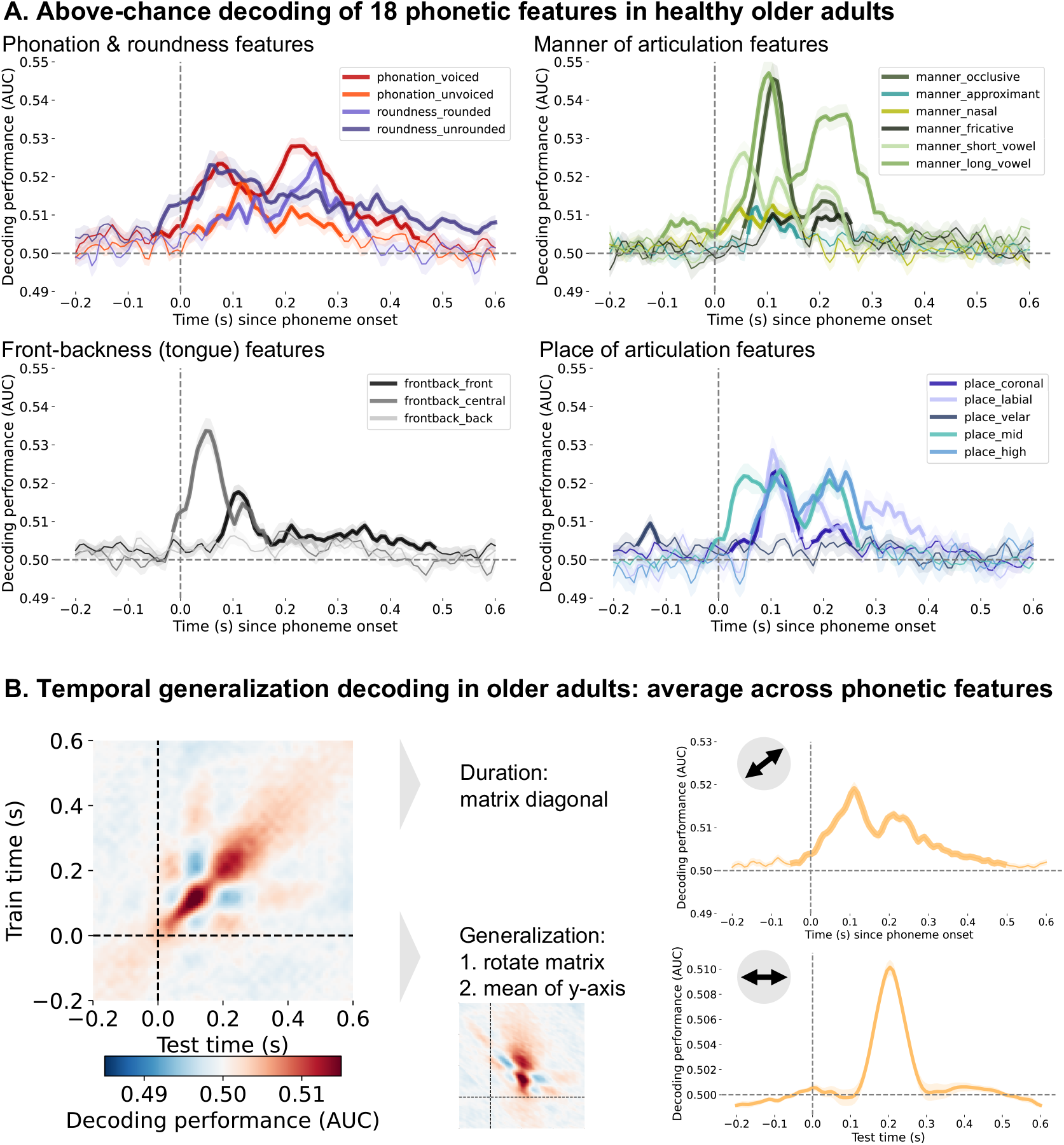
Phonetic feature decoding from EEG data in healthy older adults. **(A)** Above-chance decoding of 18 phonetic features in healthy older adults. The features vary in decoding performance and over time. The thicker line segments indicate the time points where the features are significantly decoded above chance. **(B)** The temporal generalization (TG) approach in healthy older adults revealed that phonetic features are decodable above chance for 0.5s (top right panel) and generalizable for 0.139s (bottom right panel). The decoding duration corresponds to the diagonal of the TG matrix (left panel), thus the decoding performance when the decoder was trained on the same time slice as was used for testing the decoder. To quantify the generalization time, the TG matrix is reoriented and then averaged across the y-axis.

Sustained neural encoding implies simultaneous processing of multiple phonemes. Gwilliams et al. (2022) found that the brain encodes information from the preceding three phonemes, retaining their order by encoding time since onset. Using TG analysis, they revealed that phonetic information is maintained within a neural configuration for 0.08s before evolving. This method provides insight into the speed of neural information encoding changes, demonstrating the dynamic nature of phonetic neural representation (fig.1E). Here, we investigate whether healthy older adults also show a dynamic processing scheme, given that this seems to be an important mechanism for orderly processing of phoneme sequences.

Using the TG decoding approach, we observed the train time by test time matrix shown in figure 2B (left panel), wherein each data point represents the decoding performance. The width of the significant cluster in this matrix represents the generalization time of phonetic information in a given time point and spatial configuration (i.e., a decoder trained at time t). To extract the duration of generalization, we rolled the rows of the matrix such that the values along the diagonal become oriented at a given x-coordinate. We then average across the y-axis of the matrix, resulting in figure 2B (right bottom panel). The width of the significant matrix cluster corresponds to 0.139s in healthy older adults. We compared the phonetic encoding duration (0.53s, the matrix diagonal, fig.2B right top panel) to the generalization (0.139s, the width of the reoriented matrix) using a one-sample permutation cluster test to confirm the dynamic evolution of phonetic encoding. We found a significant difference between -0.044s to 0.165s (t(23)=- 2.86, p<.001) and between 0.188s to 0.46s (t(23)=-3.12, p<.001), confirming that phonetic information is encoded dynamically, evolving over time and space, in healthy older adults. The same analysis and figures for the aphasia group can be found in the supplementary information (fig.S.2 and section S.5).

### 3.2 Phonetic encoding in individuals with aphasia

IWA have been found to have impaired phoneme identification (Kries et al., 2023) and phonological processing (Blumstein, 1998). This led us to investigate the neural dynamics of phonetic encoding in aphasia in comparison to healthy controls. Regarding the TG analysis, we expected that the aphasia group would show a different encoding pattern than the healthy control group. We tested whether IWA would show shorter encoding duration, slower speed of evolution of encoding than controls, or a mix of both (fig.1F). Figure 3A shows the TG matrices for the aphasia and control groups overlaid. We conducted a permutation cluster test to compare matrices between groups and found a cluster in which the aphasia group showed significantly lower decoding performance than the control group (p<.001; contour in fig.3A).

**Figure 3:**
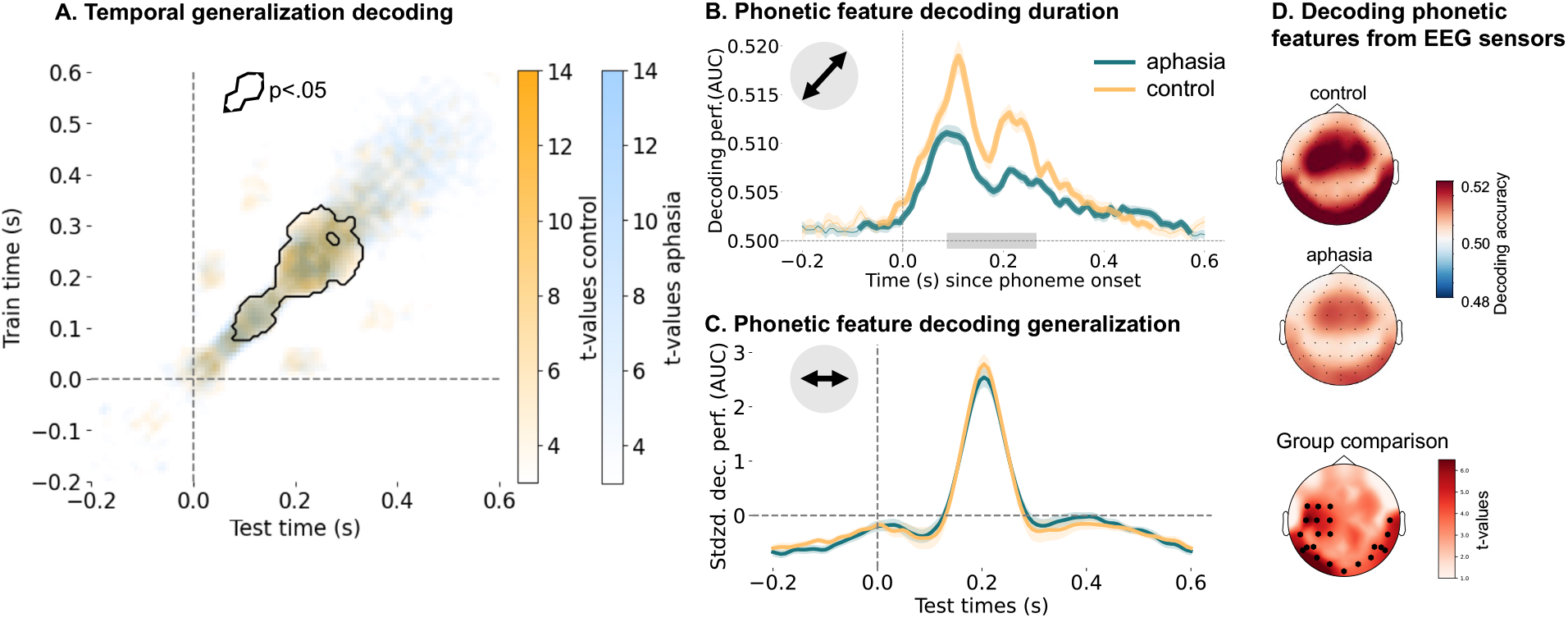
Comparing phonetic decoding between groups. **(A)** The TG matrices shown as t-values overlaid for the control (yellow) and aphasia (blue) group. To obtain this, a one sample permutation cluster test was conducted for each group separately, which calculates a t-value for each data point in the TG matrix. The black contour indicates the cluster in which the groups significantly differed from each other. To achieve this, we conducted a permutation cluster test to compare groups. **(B)** A significant difference between groups was found from 0.08 to 0.25s (as indicated by the gray bar) where the control group showed increased decoding performance. The bold parts of the lines indicate the time points where the features are significantly decoded above chance. **(C)** No significant difference in generalization was found between groups. For both groups, the encoded information was passed on to new neural ensembles every 0.14 s. **(D)** The top 2 topographies show how well phonetic features could be decoded for each of the 64 EEG sensors. The bottom topography shows the difference between both groups, specifically the t-values calculated using a mass-univariate independent samples t-test. The control group showed generally higher decoding, but especially in the 22 sensors marked in bold in the bottom topography.

The matrix diagonal shows phonetic feature decoding duration. We compared this duration between IWA and controls using a permutation cluster test. Results revealed significantly lower decoding accuracy in the aphasia group from 0.08 to 0.25s post-phoneme onset (p<.001; fig.3B, supplementary fig.S.3). Based on the two observed peaks in the decoding timecourse, we additionally investigated whether the decoding performance at each peak correlates with different behavioral measures, within the aphasia group. We hypothesized that behavioral correlates of each peak could provide insight into the associated functional processes; however, no significant correlations were found between either peak and language- or cognition-related variables (supplementary section S.8).

To compare groups on how long phonetic information is maintained within a same temporal decoder and spatial configuration, we extracted the train time lags between 0 and 0.35 s, based on previous research (Gwilliams et al., 2022), and standardized the data (fig.3A). We used a permutation cluster test to compare groups on generalization time, and found no significant difference between groups (fig.3C), with average generalization time being 0.147s in the aphasia group and 0.139s in the control group. The speed of evolution of phonetic encoding across spatial configurations in IWA is thus highly overlapping with healthy controls. Taken together, these results partially support the hypothesis in the top right panel of figure 1F: an altered duration of phonetic encoding in aphasia (i.e., weaker encoding after initial processing time), but no change to the speed with which phonetic information is passed between neural populations. However, the difference in duration of phonetic encoding that we observed is indirect, i.e., phonetic information is more weakly encoded in aphasia after initial processing time, a scenario that we did not fully capture in our hypotheses in figure 1F.

Finally, to examine the spatial underpinning of these dynamic representations, we compared sensor-level topographical differences of phonetic feature decoding between aphasia and control groups (fig.3D). With a cluster-forming p-value threshold of p<.001, we identify a cluster of 22 EEG sensors (p<.001; fig.3D) over which IWA had lower decoding accuracy than healthy controls. The sensors primarily localized to left auditory cortex and posterior sensors bilaterally.

### 3.3 Effects of lexical predictability on phonetic encoding

Given the observed difference in phonetic encoding duration between IWA and controls, we post hoc investigated potential causes. Previous research suggests lexical predictability influences phonetic encod-ing duration (Gwilliams et al., 2022). Lexical entropy, quantifying uncertainty about word identity, is typically higher for word-initial phonemes and lower for later phonemes (Gwilliams and Davis, 2022). We explored whether changes in phonetic encoding duration in aphasia relate to an altered interaction between phonetic encoding and lexical uncertainty. If so, encoding duration should correlate with lexical uncertainty: higher uncertainty would lead to longer phonetic information encoding.

We conducted two analyses to investigate this; we subset (1) all phonemes according to phoneme position relative to word onset and offset and (2) phonemes according to lexical entropy. We compared groups on each phoneme position separately using a permutation cluster test, demonstrating a significant group difference for the first (p<.001), second (p=.02) and third (p=.03) phoneme positions, as well as the second to last (p<.01) and fifth to last (p=.016) phoneme positions (fig.4A). For the first phoneme relative to word onset, we observed a difference in reactivation, wherein only the control group seemed to maintain longer or “reactivate” the phonetic features in earlier temporal decoders, i.e., 0.1 to 0.25s (supplementary fig.S.4). For the other phoneme positions, the group difference was located on or close to the TG matrix diagonal, thus displaying similar group comparison results as found when taking into account all phonemes, i.e., altered strength of encoding. We would like to point out that we might not have had enough phoneme trials beyond the second (and second to last) phoneme position (table 1) to draw strong conclusions from this analysis.

**Figure 4:**
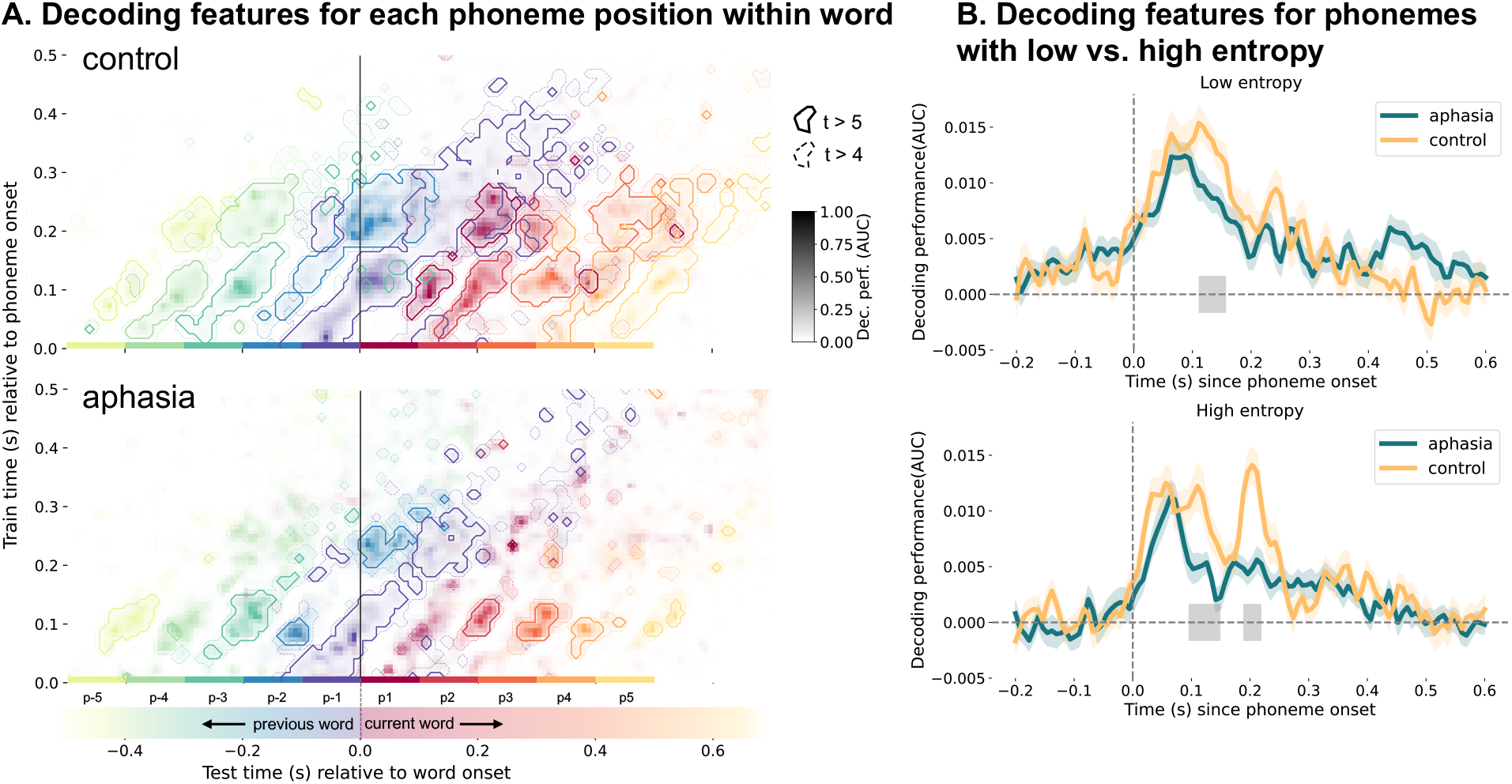
Testing the effect of phoneme position and lexical uncertainty on phonetic decoding in individuals with aphasia and controls. **(A)** The top plot shows temporal generalization (TG) decoding of phonetic features for each phoneme position within word separately. Each phoneme position is depicted in a different color. The bottom plot shows the same type of plot for the aphasia group. The TG matrices are shown as contour plots, where the contours frame t-values larger than 5 (solid lines) and 4 (dashed lines). The decoding strength is shown as gradient from light (lower decoding) to dark (higher decoding) in the respective color of each phoneme position. Phoneme positions where the decoding matrices significantly differed between groups are P1, P2, P3, P-2 and P-5. **(B)** We decoded phonetic features separately for phonemes with low and high lexical entropy. The top plot shows the difference between groups for low entropy phoneme decoding, the bottom plot shows the same graph for high entropy phoneme decoding. The gray bars indicate where the groups are significantly different from each other.

Lexical entropy is one possible way to measure linguistic prediction (Gwilliams and Davis, 2022), which is represented within the language network (Shain et al., 2020), and given that the language network is damaged in aphasia, we expect the linguistic predictions mechanism to be disrupted. Therefore, we hypothesized that IWA would show no difference in phonetic decoding performance between phonemes with low and high entropy, but that healthy controls would show such a difference.

We compared groups in low and high entropy conditions separately using permutation cluster tests. In both conditions, controls showed higher decoding than the aphasia group between 0.1s and 0.15s post-phoneme onset (low entropy: 0.110s-0.157s, average decoding: control=0.013 and aphasia=0.008, t(61)=0.757, p=0.042; high entropy: 0.095s-0.149s, average decoding: control=0.009 and aphasia=0.004, t(61)=0.002, p=0.011; fig.4B). Additionally, for phonemes with high entropy only, the control group shows also higher decoding in a window from 0.188s to 0.219s (average decoding: control=0.012 and aphasia=0.004, t(61)=1.025, p=0.002) after phoneme onset (fig.4B). Second, we tested the interaction between group and entropy condition by subtracting the low entropy condition from the high entropy condition (see supplementary fig.S.5) for each group, subsequently conducting an independent samples t-test for each time lag and finally correcting p-values for multiple comparisons (false discovery rate). Based on findings from Gwilliams et al. (2022), we tested in a time window from 0.15s to 0.35s. We found a significant interaction between group and entropy condition from 0.204s to 0.212s after phoneme onset (t(61)=2.145, p=0.037), wherein the control group had higher decoding performance than the aphasia group in the high entropy condition only. This means that phonemes with higher lexical uncertainty are encoded for longer – an adaptive encoding duration mechanism found in previous work (Gwilliams et al., 2022). We replicated this finding in healthy older adults, but find no evidence for the same processing occurring in people with aphasia.

Given the significant difference in high-entropy phoneme decoding between groups at 0.2s post-speech- onset, we explored potential behavioral correlates within people with aphasia and across all participants. We tested correlations with behavioral acoustic, linguistic, cognitive and demographic variables in an exploratory way (no correction for multiple comparisons applied), which can be found in supplementary section S.10.1. Most importantly, we found a positive correlation between the phoneme identification task performance (data available in 22 IWA and 20 controls) and high entropy phoneme decoding performance around 0.19s after phoneme onset (all participants: r=0.354, p=0.021; within-aphasia group: r=0.425, p=0.048; fig.5, supplementary section S.10.1). The phoneme identification task consisted of participants having to identify whether the sound they heard was a /bA/ or /dA/, evaluating how consistently people identify phonemes amid acoustic variability (Kries et al., 2023). This task addresses the ‘invariance problem’ — the challenge of mapping variable speech sounds to discrete phonological categories. This significant correlation occurred around the same latency that the control group showed stronger phonetic feature encoding of high-entropy phonemes than the aphasia group (fig.5).

**Figure 5:**
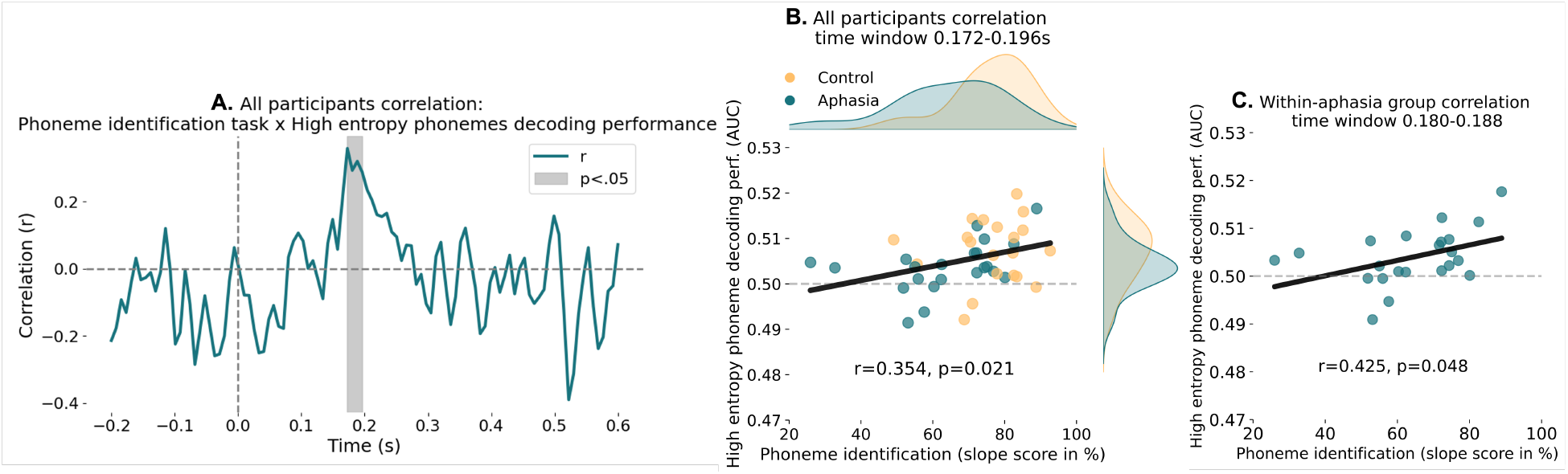
Speech sound categorization positively correlates with encoding of high-entropy phonemes around 0.2s after phoneme onset. **(A)** The high entropy phonetic feature decoding timecourse corelated with the phoneme identification task shows a significant positive correlation from 0.172 to 0.196s after phoneme onset. Here the correlation coefficient timecourse is shown across groups. **(B)** Across-groups correlation for significant time samples. Depicted are the individual data points as well as the distribution of values per group per measure. **(C)** Within-aphasia group correlation for significant time samples. Also see supplementary section S.10.1.

## 4 Discussion

In this study, we decoded phonetic features from EEG data during continuous story listening in 39 individuals with aphasia (IWA) and 24 age-matched (thus older) healthy controls. We found (1) phonetic features can be robustly decoded from 25 minutes of EEG data in healthy older adults, (2) IWA have an altered phonetic encoding duration, strength and topography compared to controls, and (3) the adaptive mechanism of phonetic encoding duration being contingent on lexical uncertainty seems to be disrupted in aphasia.

### 4.1 Replicating phonetic encoding dynamics in older adults with EEG data

Robust encoding of phonetic features was reproduced in older adults using an EEG dataset (fig.2), which replicates a prior MEG study with younger adults (Gwilliams et al., 2022). This is an important result, given that the signal-to-noise ratio is higher for MEG than EEG data (Goldenholz et al., 2009; Ahlfors et al., 2010), and given the 3.7x difference in data size between studies, i.e., 2 hours and 50,518 phonemes in Gwilliams et al. (2022) versus 25 minutes and 13,560 phonemes in the present study.

The fact that the study by Gwilliams et al. (2022) used English stimuli and here we used Dutch stimuli may also introduce variation in results due to differing phonetic features and their statistical probabilities and frequency. The different phonetic features show slightly distinct decoding timecourses, e.g., short vowels have an earlier first peak than long vowels, and phonemes with rounded lips have a later decoding peak than phonemes with unrounded lips. All phonetic features were decodable above chance at some point throughout the timecourse. Finally, we also replicated the prior finding of dynamic coding of phonetic features, in older adults. We found that the average decoding across features was above chance for 0.53 s, whereas the duration with which the same neural pattern remained informative was only 0.139 s. This suggests that in both younger and older adults, a dynamic neural pattern is used to encode and process the phonetic features of speech sounds in continuous speech.

### 4.2 Phonetic encoding in individuals with aphasia

Comparing decoding duration between groups, we found that phonetic features were decoded less well in IWA than in controls (fig.3B). Early during processing (0 to 0.08 s), no significant difference between groups was found (fig.3AB); the difference emerged from 0.08-0.25 s. IWA may initially process phonetic features, but not keep the information encoded long enough to fully integrate it with higher-level processes (Gilbert and Sigman, 2007; Sigman and Dehaene, 2005; Gwilliams et al., 2018). Thus, the duration of encoding is indirectly affected in aphasia, via lower encoding at the later processing time points.

Comparing the TG decoding performances of both groups (fig.3A and supplementary fig.S.3), it seems that the decoding performance is more diffuse, less concentrated in the aphasia than the control group. More diffuse cognitive representations in the brain have been found to be a part of the aging process, a mechanism called dedifferentiation (Koen and Rugg, 2019). Dedifferentiation, a loss of functional specialization in neural structures, is part of normal aging. While both our control and aphasia samples were older adults (average 71 years), IWA might exhibit more pronounced dedifferentiation than controls. In fact, Purcell et al. (2019) found that increased local neural response differentiation was related to less severe language impairments. This might explain the more smeared, less specific neural response to phonetic features in aphasia compared to controls. Another possibility is that the EEG data in IWA is affected by the stroke lesion site being filled with cerebro-spinal fluid, which in turn impacts the signal-to-noise ratio over sensors closest to the lesion site (Piastra et al., 2022; Salinet et al., 2014; Zbesko et al., 2018; Vorwerk et al., 2014; Cohen et al., 2015; Park et al., 2016; Cassidy et al., 2020).

Sensor-level decoding revealed similar topographical patterns across groups, but stronger encoding in controls, especially over left auditory sensors (fig.3D). This aligns with previous findings of decreased speech envelope encoding in this aphasia dataset (Kries et al., 2024). While structural changes in the left hemisphere may affect decoding accuracy (supplementary fig.S.1), functional reduction in speech processing is also possible. Current analysis cannot distinguish between structural and functional changes. A control group of right-hemisphere stroke patients without aphasia could help clarify the nature of these effects.

Given that phonological impairments have been found in IWA such as mixing up sequence order or phoneme misidentification (Blumstein, 1998), we expected to see a more square-shaped TG matrix (fig.1E), meaning that IWA would have less dynamic encoding of phonetic features and pass on the information at a slower pace between neural populations, creating more representational overlap between phonemes of a sequence. However, we instead observed that the neural pattern evolves at the same rate for both groups (fig.3). Specifically, the neural pattern completes a full evolutionary cycle every 0.14 s. This implies that phonological impairments in IWA are not related to not being able to keep apart the phonetic representations of neighbouring phonemes.

Aphasia varies widely in severity and affects different language processes (Kries et al., 2023). Group analyses may mask individual behavioral-neural relationships. Future studies should focus on more narrowly-recruited individuals with severe phonological deficits to get a better understanding of these results.

### 4.3 Effects of lexical predictability on phonetic encoding

Our third research question emerged post hoc, because we wanted to explore what might underlie changes in phonetic decoding duration/strength in aphasia. We found that at a few phoneme positions, the TG decoding matrices differed between groups (fig.4). Most markedly, the word-initial phoneme’s features – generally the phoneme with the highest uncertainty about lexical identity – were maintained longer/”reactivated” in decoders relatively early after phoneme onset in healthy controls, but not in IWA. This supports the idea that IWA may not keep phonetic features of the word-initial phoneme encoded for long enough, such that integration with higher levels of processing may be hampered. Age differences between this study (older adults) and Gwilliams et al. (20222; younger adults) may explain discrepancies in word-initial phoneme pre-activation results. Older adults might process predictive cues differently (Gillis and Kries et al., 2023), possibly reactivating and maintaining phonetic features (supplementary fig.S.4) rather than pre-activating them (Gwilliams et al., 2022). However, other methodological differences between studies limit direct comparisons. Future research could examine these hypotheses in younger and older adults.

We also assessed how lexical entropy affects phonetic encoding in IWA and controls. Controls, but not IWA, encoded phonemes with high lexical uncertainty for a longer period of time than phonemes with low lexical uncertainty. Our findings in healthy controls (older adults) align with Gwilliams et al. (2022)’s results in young adults, suggesting that the ability to flexibly adjust encoding duration based on lexical uncertainty may be crucial. IWA lack this adaptive encoding mechanism, which might thus lead to language processing impairments. IWA appear unable to encode phoneme properties for a sufficient duration to resolve lexical identity. This shows that integration between different levels of processing might be maladaptive in aphasia.

The variability in phonetic feature decoding of high-entropy phonemes around 0.2s after phoneme onset was associated with behaviorally assessed phoneme identification (fig.5, supplementary section S.10.1). This means that individuals who performed better at the phoneme identification task also showed stronger phonetic encoding when word identity was uncertain. This finding further supports that maintaining phonetic features long enough to resolve word identity is an important mechanism to support successful comprehension. Moreover, this mechanism has potential as a neurobiological marker of aphasia.

### 4.4 Applying the hierarchical dynamic coding framework to aphasia

Gwilliams et al. (2022) demonstrated that the duration for which phonetic features are maintained in the same neural population (0.08 s) match with and is contingent on the duration of the phoneme input, i.e., 0.08 s. In the current study, we found that the duration of maintaining phonetic information (0.14 s) is similar to the average phoneme duration in the stimulus (0.10 s), thus is in line with previous findings. Recently, Gwilliams et al. (2024) extended TG analysis to higher linguistic levels, revealing hierarchical dynamic coding (HDC) as a general sensory-cognitive processing principle. Given our findings of weaker, more diffuse phonetic feature encoding in IWA, this pattern might extend to other language processing levels in aphasia. If neurotypical healthy adults with intact speech comprehension show a specific coding pattern, while individuals with a language impairment show a different pattern, then maybe the neurotypical coding pattern is conditional for successful speech comprehension. It is possible that a certain spatio-temporal concentration in neural encoding of speech features (for example phonetic features) is necessary in order to efficiently integrate the information with other language representation levels.

### 4.5 Caveats

Limitations include a heterogeneous but small sample of 39 IWA. Future studies should aim to reproduce findings with larger samples, longer recordings, and extend analysis to word- or phrase-level representations. Furthermore, the lesion site and its effect on neural recordings pose a problem for more detailed interpretation of results. Therefore, it would be essential to record data from a control group that has a right-sided lesion without aphasia in the future, so as to investigate to what extent altered encoding of speech is related to structural versus functional changes in aphasia. It is generally an issue in aphasia studies that control groups most of the time consist of healthy adults, whereas a comparison with demographically-matched adults with a brain lesion without aphasia would present a more precise control group. Another limitation of the approach we employed here is that it is not reliable at an individual participant level, at least not with the relatively limited amount of data available.

### 4.6 Conclusion

Overall, we found that phonetic processing is robustly encoded in EEG responses of older adults. The primary marker of aphasia was altered strength of phonetic encoding after initial processing, rather than a slower evolution of neural representation. Moreover, phonemes with high uncertainty about word identity were encoded longer in controls than in IWA, indicating that encoding phonetic information until word identity is resolved may be a crucial mechanism for successful speech comprehension. These results suggest that speech impairments in aphasia may be driven by difficulties maintaining lower-order information long enough to recognize lexical items.

## Acknowledgements

The authors would like to thank all participants, especially all the brave participants with aphasia and their partners, family or friends that support them. Moreover, the authors would like to thank everyone who helped with data collection and recruitment: Dr. Pieter De Clercq, Dr. Klara Schevenels, Janne Segers, Rosanne Partoens, Charlotte Rommel, Dr. Ramtin Mehraram, Ines Robberechts, Laura Van Den Bergh, Anke Heremans, Frauke De Vis, Mouna Vanlommel, Naomi Pollet, Kaat Schroeven, Pia Reynaert, and Merel Dillen.

## Supplementary material

### S.1 Lesion information

MRI data collection was not a part of the planned data collection for this study, but ethical agreement was given to access the scans from the medical files of the university hospital. Thus, we had access to T2-weighted FLAIR images that were administered in the acute stage post-stroke, based on which the segmentations of the affected stroke tissue were manually performed. For 10 participants, scans were available from the chronic phase post-stroke, as they took part in an MRI study (De Clercq et al., 2024). 12 participants had participated in an earlier aphasia study, and thus we could use the already established segmentation maps.

**Figure S.1:**
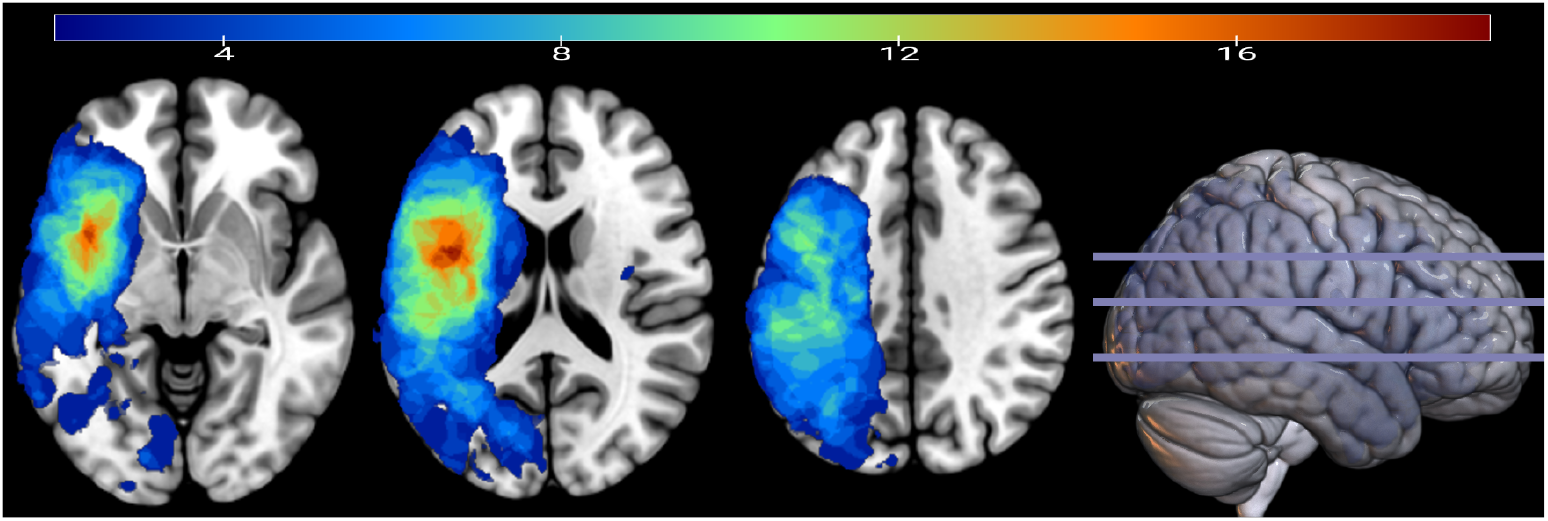
Lesion overlap image of the aphasia sample. The maximum overlap corresponds to 19 out of the total sample of 39 individuals with aphasia. Axial slices are shown in neurological orientation.

### S.2 Group demographics and behavioral variables

In supplementary table S.1, we summarized demographic information by group (details can be found in supplementary table S.2). Age, sex, education, handedness and multilinguality did not differ between groups (age: W = 464, p = 0.96; sex: *χ*^2^ = 0, df = 1, p = 1; education: *χ*^2^ = 7.26, df = 4, p = 0.1; handedness: *χ*^2^ = 0.063, df = 2, p = 0.98; multilingual: *χ*^2^ = 0.182, df = 1, p = 0.66). Details about the stroke in IWA, i.e., time since stroke onset, stroke type, occluded blood vessel, lesion location and speech-language therapy, can be found in supplementary table S.2. To visualize the damaged brain tissue of IWA, a lesion overlap image was created (supplementary fig. S.1). Demographic information was acquired via a self-reported questionnaire. Handedness was assessed via the Edinburgh Handedness Inventory (Oldfield, 1971). All behaviorally assessed tests are described in detail in supplementary section S.4.

**Table S.1:** Demographics, language-diagnostic information and covariates by group.

|  |  | Control | Aphasia |  |
| --- | --- | --- | --- | --- |
| **Demographics** |  |  |  |  |
| Age in years (mean(SD))<sup>◊</sup> |  | 71.5(7) | 69.5(12.4) | n.s. |
| Sex (n(%)) | female | 8(33.3%) | 13(33.3%) | n.s. |
|  | male | 16(66.7%) | 26(66.7%) |  |
| Education (n(%)) | primary education | 0(0%) | 3(7.7%) | n.s. |
|  | secondary education | 5(20.8%) | 15(38.4%) |  |
|  | tertiary education/college | 8(33.3%) | 10(25.6%) |  |
|  | Master's degree | 9(37.5%) | 11(28.2%) |  |
|  | PhD | 2(8.3%) | 0(0%) |  |
| Handedness (n(%)) | right | 22(91.6%) | 35(89.7%) | n.s. |
|  | left | 1(4.2%) | 2(5.1%) |  |
|  | ambidextrous | 1(4.2%) | 2(5.1%) |  |
| Multilingual (n(%)) | yes | 22(91.7%) | 33(84.6%) | n.s. |
|  | no | 2(8.3%) | 6(15.3%) |  |
| **Language diagnostic tests** |  |  |  |  |
| ScreeLing (max=72) (mean(SD)) |  | 69.9(2.5) | 61.5(9.4) | * * * |
| Picture naming (max=276) (mean(SD)) |  | 271.3(4.1) | 224.7(58.1) | * * * |
| **Covariates** |  |  |  |  |
| Hearing (Fletcher index in dB HL) (mean(SD)) |  | 24.5(12) | 28.4(14.1) | n.s. |
| Cognition (OCS composite score in %) (mean(SD)) |  | 94.2(4.7) | 80.4(16.6) | * * * |
| Alertness (Likert scale) (median(range)) |  | 4.75(3.5-5) | 3.5(2-5) | ** |
| Fatigue (Likert scale) (median(range)) |  | 1.5(1-4.5) | 3.5(1-4.5) | ** |

<sup>◊</sup> Age is also used as a covariate; n.s. = not significantly different; p<0.05\*; p<0.01\*\*; p<0.001\*\*\*

### S.3 Demographic and lesion information of individuals with aphasia

**Table S.2:**
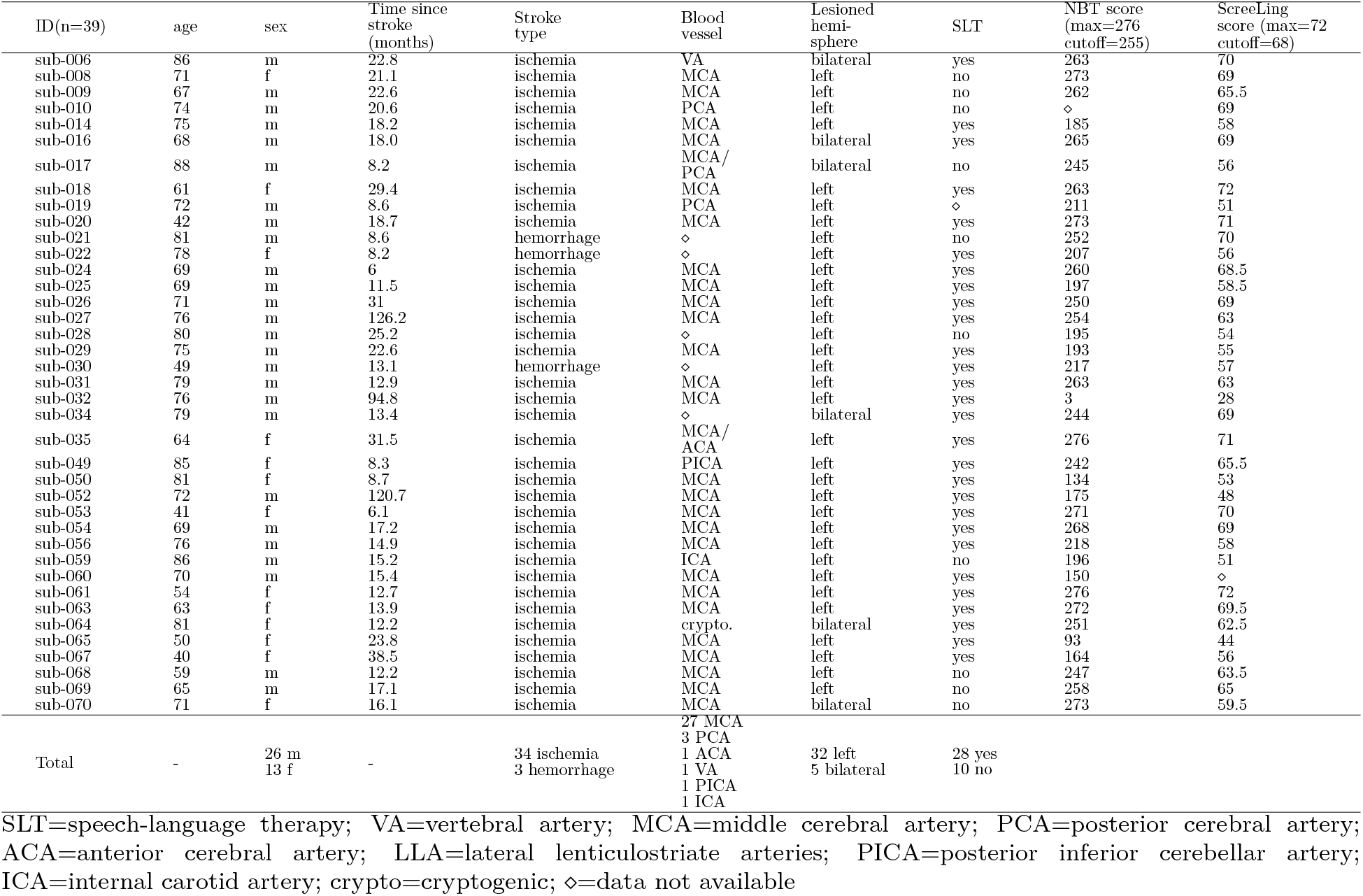
Demographic, lesion and diagnostic information of the aphasia group.

### S.4 Behavioral assessment

Behaviorally assessed measures: alertness, fatigue, hearing, rise time discrimination, phoneme identification, ScreeLing (phonology, semantics, syntax, general reaction time), picture naming, phonological word fluency, semantic word fluency, cognition (attention, memory, executive function, hemispatial neglect), story comprehension questions.

#### S.4.1 Alertness and fatigue ratings

Subjective ratings of alertness and fatigue were administered at 3 time points throughout the data collection session, hereafter referred to as t1, t2 and t3. The EEG measurement took place between t1 and t2, therefore we averaged across t1 and t2 of the alertness and fatigue rating respectively. Individuals with aphasia reported to be significantly less alert and more fatigued than controls on this average score (see results in table S.1). T3 took place roughly 1 hours after t2, in this hour between t2 and t3, participants did cognitive and language tests.

We also analyzed the changes over time points of alertness and fatigue ratings between groups. Linear mixed effects models were employed for this purpose. The analysis of alertness revealed a main effect of group (p<0.001), but no main effect of time point or interaction effect (time point: p=0.06; interaction: p=0.226). Thus, across time points, individuals with aphasia were less alert than healthy controls. The analysis of fatigue showed significant main effects (group: p<0.001; time point: p=0.015) and a significant interaction effect between group and time point (p=0.02). Post hoc testing revealed that while the initial level of fatigue was at a similar level for both groups at the beginning of the test session (t1; p=0.92), individuals with aphasia were more tired than healthy controls after the first part of the experimental session (t2; p<0.001) and at the end of the protocol (t3; p=0.002).

We offered IWA to continue the administration of the second part of the experimental protocol (everything after t2) on another day in case they felt too exhausted after the EEG measurement. Indeed, 5 out of 39 IWA felt too exhausted at t2 and chose to do the rest of the tests on another day. This second session took place at the patients’ homes, so that they did not have to arrange transportation to the lab again.

#### S.4.2 Hearing

Hearing thresholds were assessed via pure tone audiometry at frequencies ranging from 0.25 to 4 kHz. In case the hearing thresholds below 4 kHz were *>*25 dB HL, this information was used to increase the amplitude of stimulus presentation during the EEG measurement. The pure tone audiometry thresholds at 0.25, 0.5 and 1 kHz were averaged and then divided in half to come to the amount of dB that was added to the stimulus presentation amplitude of 60 dB SPL during the EEG paradigm. This calculation was done for each ear separately. After a short example stimulus, participants were asked whether the loudness was comfortable and if necessary the presentation volume was adjusted. The degree of volume adjustment did not differ between groups (W = 170, p = 0.8). The Fletcher index (average of hearing thresholds at 0.5, 1 and 2 kHz) was calculated per ear and subsequently averaged across both ears. The Fletcher index did not differ between IWA and healthy controls (W = 541.5, p = 0.29, confidence interval: [-2.49 10]; table S.1).

#### S.4.3 Acoustic processing

The rise time discrimination task measures how well participants discriminate the rate of change in amplitude at the onset of a sound. Precisely, the task was presented as a three-alternative forced choice task, where the deviant stimulus had to be discriminated from two identical reference stimuli. The stimuli were created in MATLAB using one-octave noise bands centered at 1 kHz (Van Hirtum et al., 2019). Stimuli were calibrated and presented in the left ear at 70 dB SPL (sound pressure level). The reference stimulus had a rise time of 15ms. The deviant stimuli were computed to have rise times that decreased logarithmically in 50 steps from 699ms to 16ms. The duration of each stimulus was 800ms. The number of trials differed between participants, as the task followed a one-up/two-down adaptive staircase procedure. This means that after two correct responses in a row, the difference in rise time between stimuli became smaller, thus more difficult, during the next trial. After one erroneous response, the difference in rise time between stimuli became larger, thus easier to discriminate. This way, a threshold corresponding to 70.7% correct was targeted (Levitt et al., 1971). The task ended once 8 reversals (i.e., changes in direction) were reached. In case no reversals were present, the task ended after a maximum of 87 trials. The rise times of the deviant stimulus of the last 4 reversals were averaged to determine the final threshold. The aphasia group performed significantly lower than the control group on this task (F=23.29, p<.001).

To make sure that the task was well understood by all participants, they performed between 4 and 8 practice trials before starting the task.

#### S.4.4 Phoneme processing

The phoneme identification task assesses how consistently speech sounds (here /bA/-/dA/) are identified. We used the same task and stimuli as employed in Vandermosten et al. (2010). The task was presented as a two-alternative forced choice identification task. Participants were instructed to decide whether the stimulus they heard sounded more like a /bA/ or more like a /dA/. The stimuli were created based on a naturally spoken /bA/. The first 100 ms of the second formant (F2) of this syllable was linearly interpolated in 10 steps to create the stimuli, using Praat (Praat (Boersma and Weenink, 2022); see Vandermosten et al., 2010 for more details). The difference between /bA/ and /dA/ solely relies on the F2 slope, this way a gradual continuum was created between these speech sounds. Thus, distinguishing between the two speech sounds relies mostly on dynamic cues, namely the discrimination of the spectral changes of F2 over time (i.e., whether the F2 slope is rising or falling). During the task, each of the 10 stimulus steps was presented 8 times in a randomized order, i.e., 80 trials. The stimuli were calibrated and presented monaurally at 70 dB SPL. At the start of the task, the two speech sounds at the extremities of the stimulus spectrum were presented as reference practice trials, to make sure that the task was well understood by all participants.

To score this task, the amount of /dA/ responses for each stimulus step was taken and divided by 8 (i.e., number of presentations per stimulus step), to arrive at the proportion of /dA/ responses. This allowed us to fit a psychometric curve on the data points using the toolbox Psignifit in MATLAB (https://github.com/wichmann-lab/psignifit). This toolbox allows to fit subject-specific guess and lapse rates, thereby we avoided making assumptions about performance at the extremities of the stimulus continuum, hence the slope was not affected by such assumptions. As borders for the guess rate, we defined a range between 0 and 0.89 and for the lapse rate a range between 0 and 0.1 on the scale of proportion of /dA/ responses. We used uniformly distributed priors in order to avoid biasing the definition of the lapse and guess rate. Subsequently, the slope at the subject-individual 50% point was computed using the function getSlope from the same toolbox and was used for statistical analyses. It is an indicator of how consistently participants were able to categorize the stimulus steps, which is indicated by the steepness of the slope. In the aphasia group, 36 IWA completed the phoneme identification task, while 3 IWA experienced the task as too difficult after some initial trials. All 24 healthy controls completed the task. As a quality check, the confidence intervals of the lapse and guess rates were analyzed. Participants whose confidence interval of either of the asymptotes included 0.5 on the y-axis (i.e., the proportion of /dA/ responses) were excluded from the analysis, i.e., 14 IWA and 4 healthy controls. Thus, all statistical analyses involving this test were performed on 22 IWA and 20 healthy controls. IWA had a decreased performance on this task compared to the control group (t=-3.02, p=0.004).

An individual deviance analysis, as described in previous literature (Ramus et al., 2003; Law et al., 2014; Boets et al., 2007), was performed on the phoneme identification task. In essence, for this analysis a reference distribution was created based on a trimmed control group, and IWA were considered to deviate from this norm when their score exceeded 1.65 standard deviation (SD). More specifically, in a first step the lowest performing 5% of the control group were removed from the control group, which will be referred to as trimmed control sample. The mean and SD of the trimmed control sample were then used to standardize the raw task scores of all participants (IWA and all healthy controls). The deviance threshold was then defined at -1.65 SD of the z-scored distribution for the phoneme identification task. Scores below the deviance threshold were viewed as deviant from the control sample. We found that 11/22 IWA (50%) had deviant or impaired phoneme identification performance, while 2/20 controls (10%) showed impaired performance.

#### S.4.5 Language assessment

**ScreeLing** (Visch-Brink et al., 2010): This test consists of 3 subtests, phonology, semantics and syntax, and adds up to maximum 72 points, with the clinical cut-off threshold being 68 points. 25/39 IWA (64%) scored below cu-toff threshold and 4/24 controls (16.6%) scored below cu-toff threshold. The aphasia group had significantly lower scores than the control group at this task (see table S.1). We decided to assess this task on a tablet, such that we can get participants’ reaction times. IWA had slower reaction times than controls (F=15.48, p<.001).

The phonology subtest consists of four tasks, i.e., spoken word repetition, reading out loud, minimal pair discrimination (e.g., ‘vase’, ‘base’; Were the two words you heard identical?) and initial phoneme identification (e.g., you will hear a word and have to choose which letter is the first one of that word). Each of the tasks consists of 6 items, hence the total score on the phonology subtest is 24. The cut-off score on this subtest is at 23 (Jiskoot et al., 2023). 28 out of 39 (72%) individuals with aphasia scored below the clinical cut-off threshold. The aphasia group scored significantly lower than the control group on this subtest (t=-5.33, p< 0.001).

The semantic subtest consists of four tasks, i.e., word-picture matching (e.g., which of these 6 pictures does the word ‘cup’ fit to?), semantic congruence judgment in sentences (e.g., is this a correct thing to say, yes or no? ‘Did you already fry the wine?’), word-word matching (e.g., does ‘tree’ fit best with ‘freezer’, ‘shrub’, ‘grain’, or ‘bouquet’?), semantic pattern categorization (e.g., which word differs from the other 3 words? ‘violin’, ‘siren’, ‘trumpet’, ‘piano’). The total score on the semantic subtest is 24. The cut-off score on this subtest is at 23 (Jiskoot et al., 2023). The aphasia group scored significantly lower than the control group on this subtest (t=-3.57, p< 0.001).

The syntax subtest consists of four tasks, i.e., verb/sentence-picture matching (e.g., does the sentence ‘he is biking’ fit best with i) a picture of a man biking, ii) a picture of a bike, or iii) a picture of a man standing next to a bike?), non-canonical sentence comprehension, grammatical judgment in sentences (e.g., is this a correct sentence? ‘These flowers is too expensive.’), sentence completion (e.g., fill in the missing word: ‘… dreams of a world record’. Is it ‘she’, ‘we’, ‘his’ or ‘their’?). The total score on the syntax subtest is 24. The cut-off score on this subtest is at 23 (Jiskoot et al., 2023). The aphasia group scored significantly lower than the control group on this subtest (t=-4.19, p< 0.001).

**Picture naming test** (Van Ewijk et al., 2020): The Dutch picture naming test involves showing participants pictures of objects. The Nederlandse Benoemtest (Dutch naming test) (NBT) consists of 92 items (pictures) that vary in low, middle and high age of acquisition and word frequency. The maximum score in the test is 276, while the clinical cut-off score is at 255 points. 23/39 IWA (59%) scored below cu-toff threshold and 0/24 controls (0%) scored below cu-toff threshold. The aphasia group had significantly lower scores than the control group at this task (see table S.1).

#### S.4.6 Cognitive performance

##### Phonological word fluency

We administered the phonological word fluency subscale of the Comprehensive Aphasia Test (Swinburn et al., 2004). Participants were asked to enunciate as many words as possible that start with the letter ‘s’ within one minute. The score consists of the number of correct words expressed. Phonological word fluency tasks require recruitment of linguistic functions, such as phonological processing and knowledge. However, note that phonological fluency tasks also involve cognitive functions, such as attention, executive functions and memory. The aphasia group elicited significantly less words than the control group (F=19.7, p<.001).

##### Semantic word fluency

We administered the semantic word fluency subscale of the Comprehensive Aphasia Test (Swinburn et al., 2004). The semantic word fluency test consists of listing as many words as possible that belong in a semantic category (e.g., animals) within 1 minute. It evaluates executive functions as well as semantic representations and word retrieval (Ruff et al., 1997). The score consists of the number of correct words expressed. The aphasia group elicited significantly less words than the control group (F=39.24, p<.001).

##### Oxford Cognitive Screen-NL

(Huygelier et al., 2019): The Oxford Cognitive Screen-NL was administered to assess cognitive functioning. This test was designed to be language-independent, such that cognitive functioning can be disentangled from language functioning, which is especially important for IWA. Due to limited time in the testing protocol, we chose to only assess 4/10 subscales, i.e. attention and hemispatial neglect, reading, executive functioning, and memory. *Hemispatial neglect* was used as a means to potentially exclude participants in case they had too severe hemispatial neglect, which could bias outcomes at most of the administered tests. However, the highest hemispatial neglect score was still at a very mild level and thus we decided to not exclude any participants based on hemispatial neglect.

The task to assess *attention* consisted of crossing out target shapes among distractor shapes. The task to assess *executive functions* consisted of connecting circles and triangles in alternation in descending order of size. The *memory* task consisted of free recall and recognition of words (from the sentence read for the reading task) and shapes. These 3 tasks were used to calculate a composite score of cognitive functioning. This score was calculated by transforming the raw scores of each test into percentages and then averaging across the three outcomes. The composite score was used to regress out differences in cognitive functioning to explore neural tracking differences between groups. The cognition composite score was significantly lower in IWA than in healthy controls (W = 209.5, p < 0.001). Since the memory task was language-dependent, we also assessed the cognition composite score of attention and executive functions only and found that participants with aphasia had significantly lower cognitive scores than the control group independent of language impairments.

#### S.4.7 Story comprehension question during EEG paradigm

The story *De wilde zwanen* (The Wild Swans), written by Hans Christian Andersen and narrated by a female Flemish-native speaker, was cut into 5 parts of on average 4.84 minutes (SD: 9.58 seconds) each. The story was presented bilaterally via shielded ER-3A insert earphones (Etymotic Research) at an amplitude of 60 dB SPL (A weighted), except if hearing thresholds were above 25 dB HL at the pure tone audiometry, in which case the presentation volume was augmented (see section S.4.2). After each story part, participants answered a yes/no question and a multiple choice question about the content of the preceding story part. As these questions were not validated, we did not assess them. They were solely introduced in the protocol to make participants follow the content of the story attentively. For both types of questions, we calculated an accuracy score and found a significant group difference (Yes/No question: W = 300, p = 0.01; Multiple choice question: W = 180, p < 0.0001).

### S.5 Phonetic feature decoding duration in aphasia

We assessed significance of the decoding time course of each of the 18 phonetic features using a one-sample cluster permutation test across subjects. We found that all 18 phonetic features were decodable above chance in the aphasia group (fig.S.2A). For above-chance decoding moments, the following are the time window, average decoding performance, average t-value and p-value per feature (also see fig.S.2A):

- voiced: time window=-0.06 to 0.6s, mean AUC=0.508, mean t-value=1.415, p<0.001
- unvoiced: time window=0.04 to 0.18s, mean AUC=0.507, mean t-value=-1.120, p<0.001
- nasal: time window=0.009 to 0.149s, mean AUC=0.506, mean t-value=1.088, p<0.001
- fricative: time window=0.040 to 0.172s, mean AUC=0.506, mean t-value=0.391, p<0.001
- occlusive: time window=-0.005 to 0.304s, mean AUC=0.509, mean t-value=-0.119, p<0.001
- approximant: time window=0.064 to 0.133s, mean AUC=0.505, mean t-value=0.273, p=0.003
- approximant: time window=0.196 to 0.250s, mean AUC=0.504, mean t-value=1.459, p=0.009
- approximant: time window=0.429 to 0.483s, mean AUC=0.504, mean t-value=1.180, p=0.009
- short vowel: time window=0.017 to 0.250s, mean AUC=0.508, mean t-value=-0.509, p<0.001
- long vowel: time window=0.025 to 0.335s, mean AUC=0.514, mean t-value=1.125, p<0.001
- coronal: time window=0.017 to 0.266s, mean AUC=0.506, mean t-value=0.831, p<0.001
- labial: time window=0.056 to 0.172s, mean AUC=0.510, mean t-value=0.972, p<0.001
- labial: time window=0.266 to 0.374s, mean AUC=0.508, mean t-value=0.598, p<0.001
- velar: time window=0.188 to 0.242s, mean AUC=0.504, mean t-value=3.079, p=0.005
- mid: time window=0.009 to 0.297s, mean AUC=0.510, mean t-value=-0.419, p<0.001
- high: time window=0.087 to 0.141s, mean AUC=0.507, mean t-value=-0.957, p<0.001
- rounded: time window=0.056 to 0.328s, mean AUC=0.507, mean t-value=0.886, p<0.001
- rounded: time window=0.343 to 0.421s, mean AUC=0.505, mean t-value=1.078, p=0.002
- unrounded: time window=-0.2 to -0.114s, mean AUC=0.505, mean t-value=3.147, p=0.008
- unrounded: time window=-0.067 to 0.6s, mean AUC=0.509, mean t-value=2.474, p<0.001
- front: time window=0.048 to 0.172s, mean AUC=0.507, mean t-value=2.376, p<0.001
- front: time window=0.196 to 0.506s, mean AUC=0.503, mean t-value=1.847, p<0.001
- central: time window=-0.005 to 0.172s, mean AUC=0.508, mean t-value=0.405, p<0.001
- back: time window=0.056 to 0.172s, mean AUC=0.504, mean t-value=2.385, p<0.001
- back: time window=0.188 to 0.297s, mean AUC=0.504, mean t-value=2.633, p<0.001

When averaging across phonetic features, we observed above chance decoding from -0.08 to 0.56s (p<.001) relative to phoneme onset for the aphasia group (fig.S.2B).

**Figure S.2:**
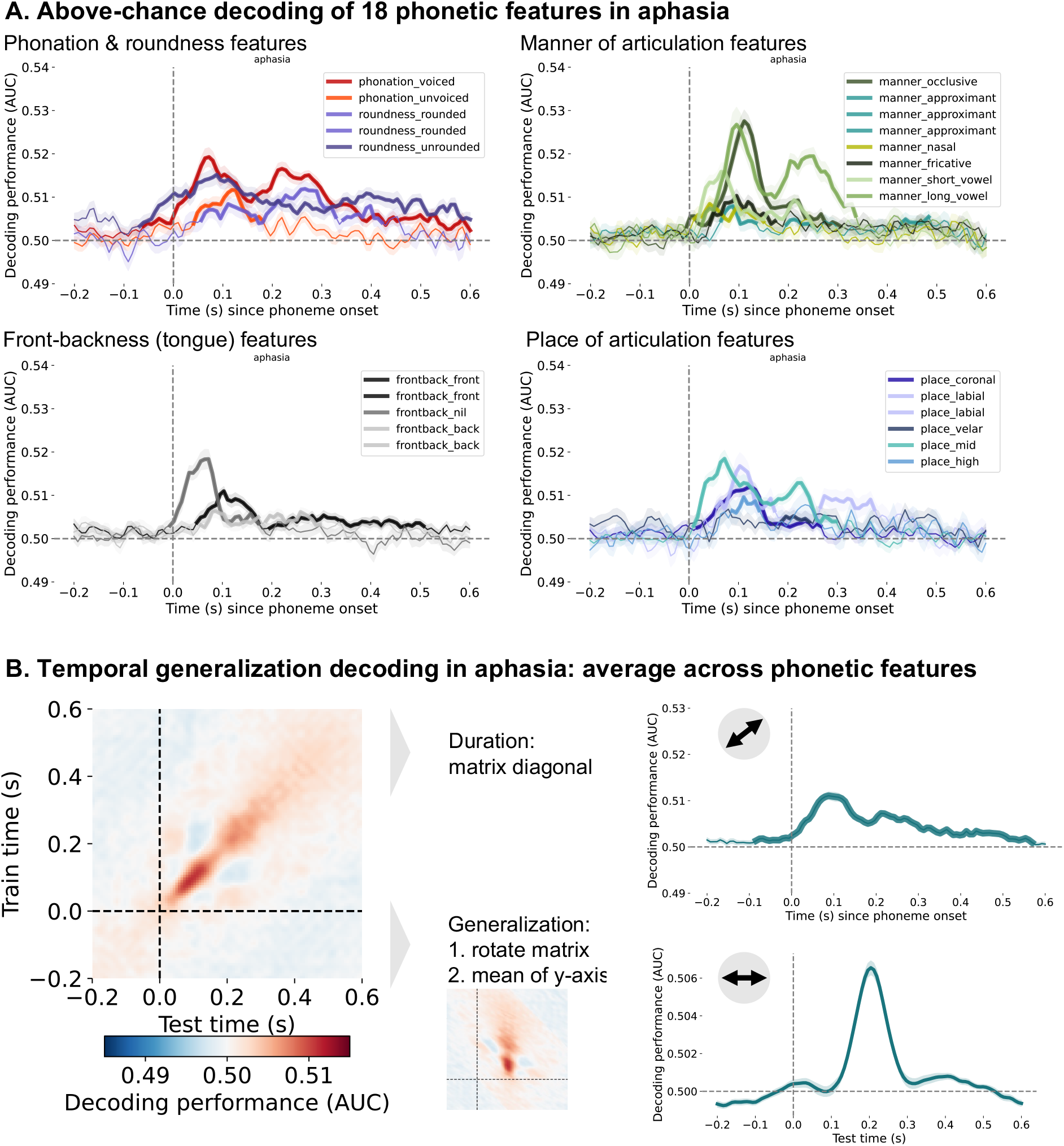
**A**. Above-chance decoding of 18 phonetic features in aphasia. The features vary in decoding performance and over time. The bold parts of the lines indicate the time points where the features are significantly decoded above chance. **B**. The temporal generalization (TG) approach in aphasia revealed that phonetic features are decodable above chance for 0.6s (top right panel) and generalizable for 0.13s (bottom right panel). The decoding duration corresponds to the diagonal of the TG matrix, thus the decoding performance when the decoder was trained on the same time slice as was used for testing the decoder. To quantify the generalization time, the TG matrix is rotated and then averaged across the y-axis.

### S.6 A more detailed view of the TG matrix group comparison

**Figure S.3:**
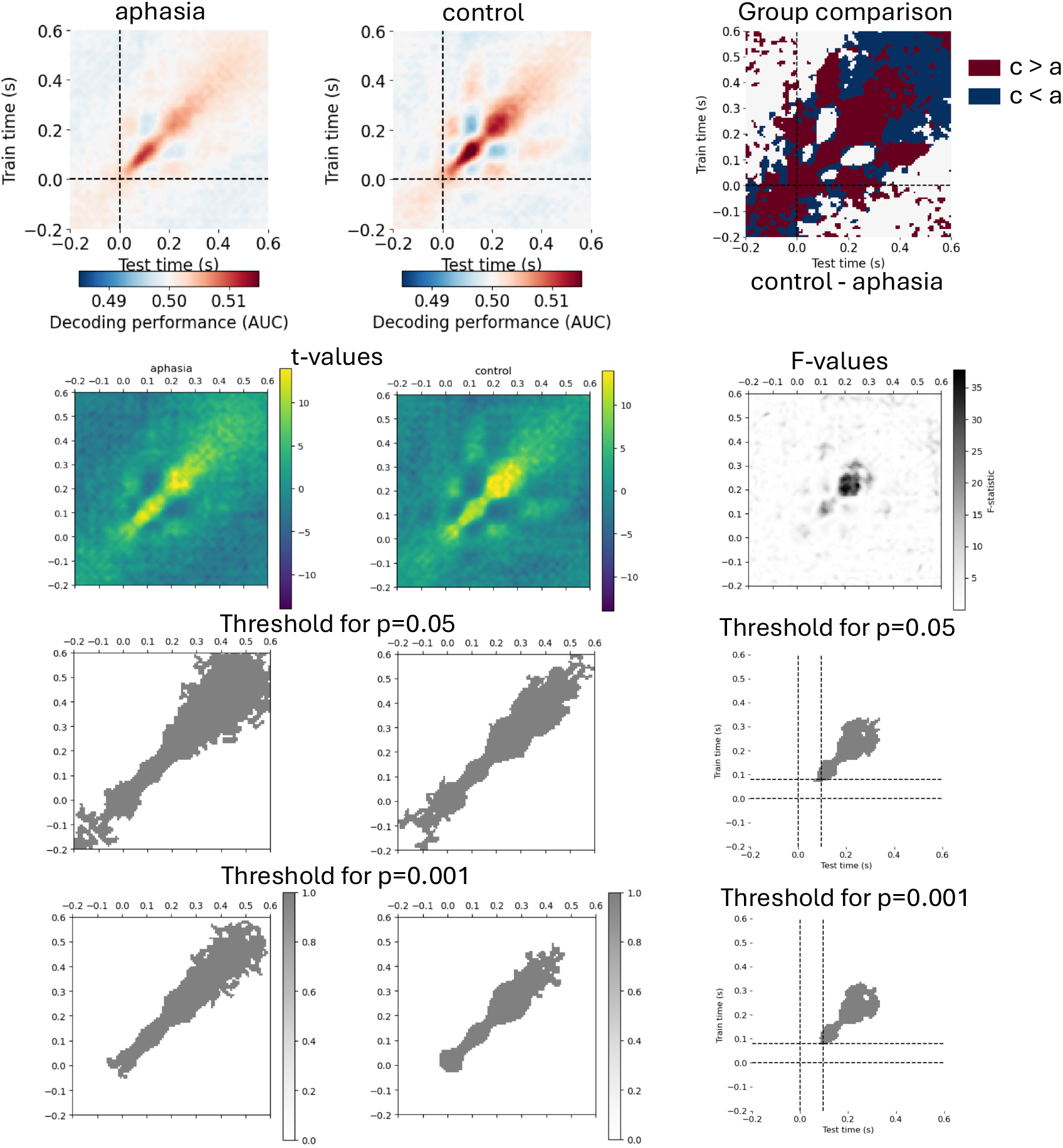
Average across phonetic features decoding in aging and aphasia. **First row:** decoding performance per group, right side panel: aphasia (a) group subtracted from control (c) group. **Second row:** (one-sample) permutation cluster tests’ t-values/F-values. **Third row:** significant clusters for each group and between groups for a cluster-forming p-value threshold of 0.05. **Fourth row:** significant clusters for each group and between groups for a cluster-forming p-value threshold of 0.001

### S.7 Duration of above-chance phonetic feature decoding in aphasia and controls

In addition to directly comparing control and aphasia groups, we investigated for how long phonetic features are decodable above chance-level in both groups separately. We used a one-sample permutation cluster test for each group separately.

As indicated in section 3.1, in **controls** we observed above chance decoding from -0.04 to 0.49 s (p<.001) relative to phoneme onset (fig.2B top right plot). Thus, on average, phonetic features are decodable for a span of 0.53 s in the control group. In the **aphasia** group, we observed significant above-chance decoding from -0.083 to 0.568 s, thus a span of 0.65 s (fig.3B bold blue line).

In a second step, we compared the above-chance phonetic feature decodability duration between aphasia and control groups. To do so, we 1) extracted the TG matrix diagonal for each individual participant, 2) calculated the standard deviation across the individual time course to compute a confidence interval for each value in the time course, 3) summed up how many time course values’ confidence intervals did not contain zero, giving us the amount of significant time lags per participant and 4) conducted an independent samples t-test with 10k permutations to compare the number of significant time lags between the aphasia and control group. This way, we found that the duration of decodability in the aphasia group is not significantly longer than in the control group (t=-0.066; p=0.9).

### S.8 Is the phonetic decoding performance at different peaks associated with behavioral measures within the aphasia group?

We hypothesized that the decoding performance in window 1 would be related to different language processes than the decoding performance in window 2. However, the language-diagnostic, cognitive and anatomical variables that we were primarily interested in did not significantly correlate with the decoding performance in either window. Only age significantly correlated with decoding performance in window 1, specifically the older individuals with aphasia are, the higher the decoding performance was from 0.05s to 0.15s after phoneme onset (r=0.32, p<0.05).

### S.9 Phoneme position analysis

**Figure S.4:**
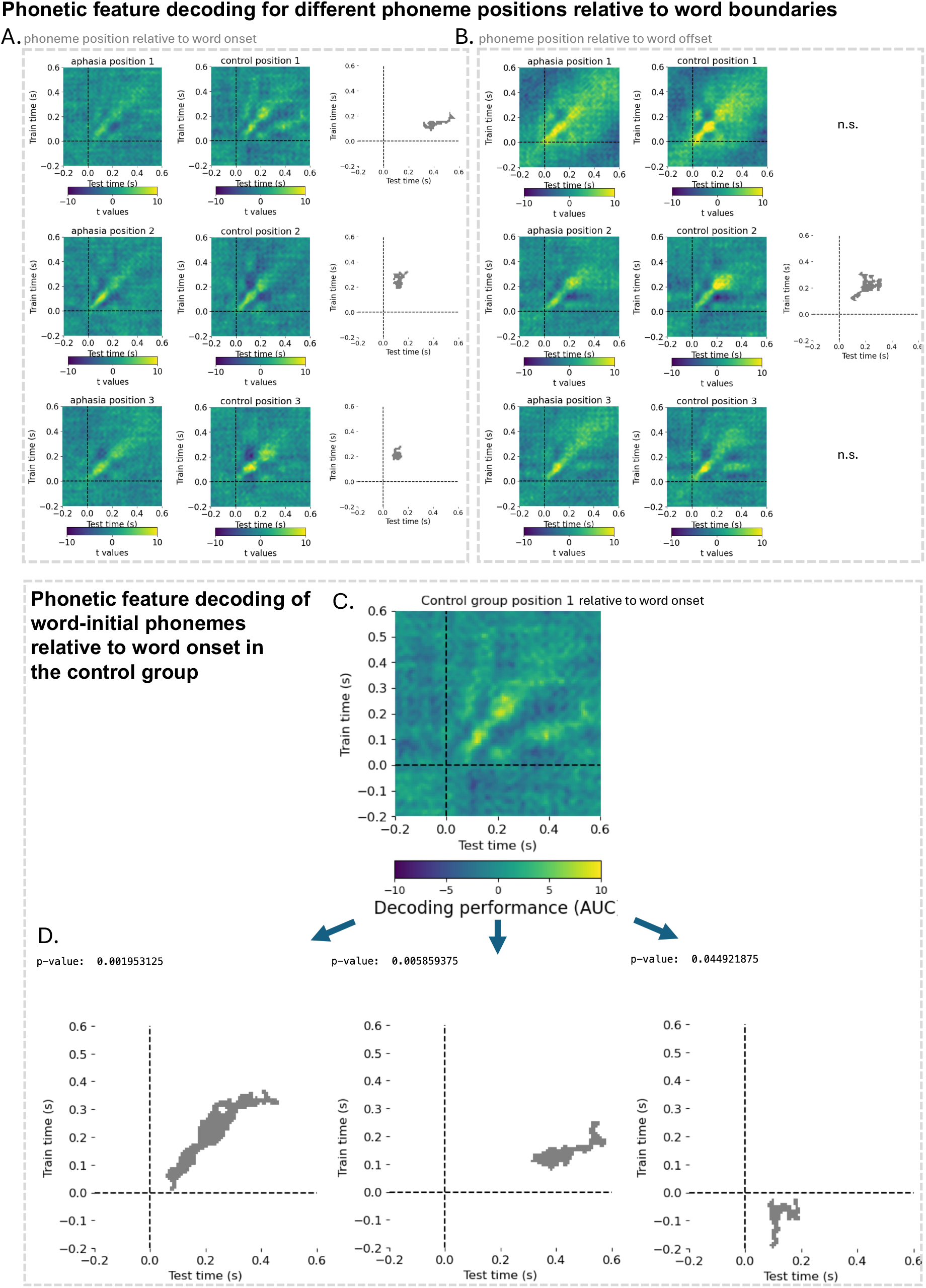
Upper panels: Phonetic feature decoding for different phoneme positions relative to word boundaries. **(A)** TG decoding matrix for first, second and third phoneme positions relative to word onset. **(B)** TG decoding matrix for first, second and third phoneme positions relative to word offset. For both panels, the first column shows decoding for the aphasia group, the second column for the control group and third column shows the cluster that was significantly different between groups. **Lower panels: Phonetic feature decoding of word-initial phonemes relative to word onset in control group**. To see whether healthy older adults have a similar pre-activation of phonetic features as found in younger adults in a previous study (Gwilliams et al., 2022), we tested the word-initial phoneme position’s feature decoding of the control group (**panel C**) against chance. We observed 3 significant clusters (**panel D**), but none of them appears to correspond to a pre-activation, thus above the diagonal. We were not able to replicate a pre-activation of phonetic features of the word-initial phoneme in healthy older adults. n.s.=not significant

### S.10 Entropy analysis

**Figure S.5:**
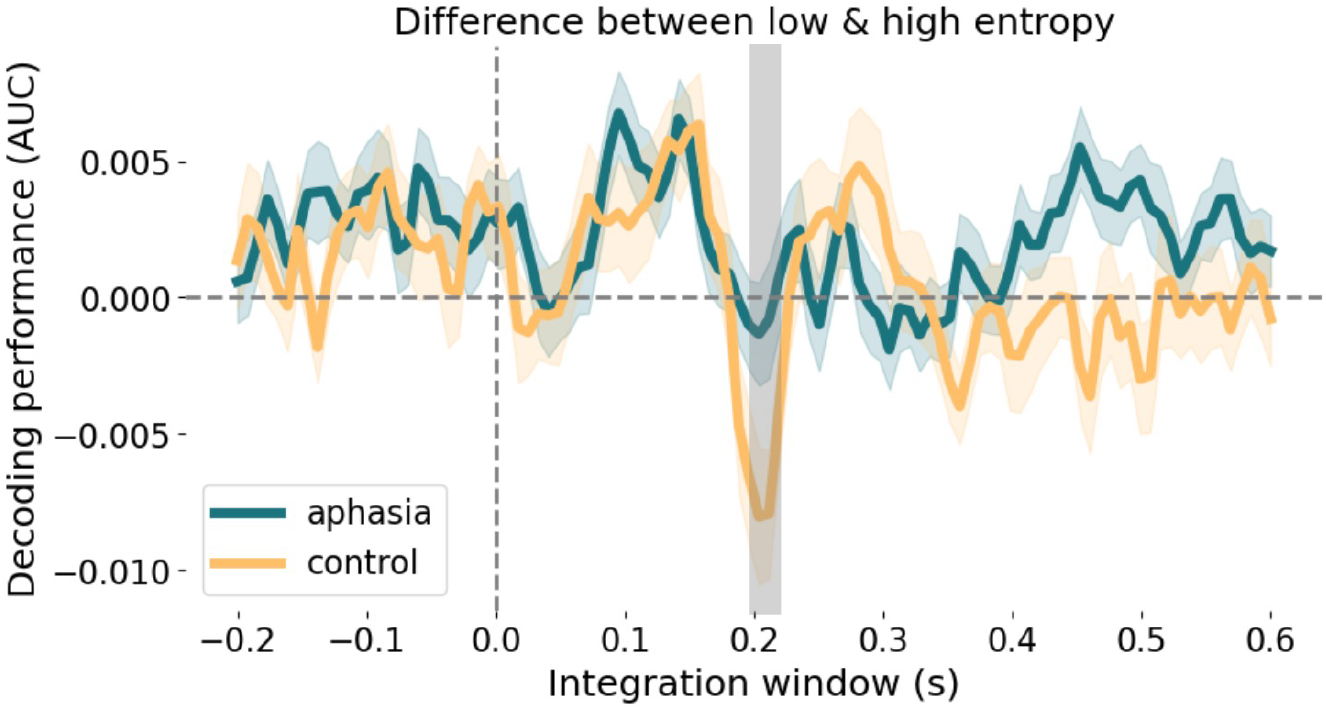
Difference between phonetic feature decoding performance of low and high entropy. The grey bar indicates time lags where the groups significantly differ.

#### S.10.1 Associations between high entropy phoneme encoding and behavioral variables

We tested in an exploratory way whether the high entropy phoneme decoding timecourse correlates with behavioral measures within the aphasia group. As this is an exploratory analysis, we did not do any corrections for multiple comparisons.

Most importantly, we observed that the better individuals with aphasia were at the phoneme identification task (i.e., consistently identifying phonemes, regardless of acoustic variance), the stronger they encoded phonetic features of high-uncertainty phonemes around 0.19s after phoneme onset (fig.5C). This was also true when we tested for correlations across groups (fig.5A and B).

The rise time discrimination task positively correlated with high-uncertainty phoneme encoding around 0.02s (r=0.3) and 0.22s (r=0.4) after phoneme onset. The score on the rise time discrimination task corresponds to the smallest difference between target and standard stimuli that people could perceive, meaning that lower values indicate better performance at this task. Thus, a positive correlation means that the bet-ter the rise time discrimination performance is, the weaker phonetic features of high-uncertainty phonemes are encoded. A negative correlation between rise time discrimination performance and phoneme encoding was found around 0.38s after phoneme onset (r=-0.4). Thus here, the better the rise time discrimination performance is, the stronger phonetic features of high-uncertainty phonemes are encoded.

Tests that assessed general language impairment severity (i.e., picture naming test, ScreeLing test, yes/no question accuracy and multiple-choice question accuracy) showed a negative correlation around 0.15 s after phoneme onset. This means the worse the language performance was, the stronger they encoded phonetic features of high-uncertainty phonemes at this point in the decoding timecourse.

### S.11 Participant recruitment

Between October 2018 and April 2022 (with a COVID-19-related break between March and June 2020), patients were recruited via daily screening at the stroke unit of the university hospital Leuven (score ≤ cut-off threshold on the Language Screening Test (LAST), Flamand-Roze et al., 2011) or via advertising the study in speech-language pathologists’ practices and rehabilitation centra (patients with a formal aphasia diagnosis). During that period (October 2018 and April 2022), 1782 patients were admitted to the stroke unit. Of those, 1010 were screened with the LAST. For data collection in the chronic phase post-stroke, we only included IWA that had no formal diagnosis of a psychiatric or neurodegenerative disorder and that had a left-hemispheric or bilateral lesion. 148 individuals were eligible to participate in the chronic phase and were contacted. 38 patients agreed to participate, and further 7 individuals with aphasia were recruited via flyer advertising, totaling EEG and behavioral data of 45 individuals with aphasia. Unfortunately, 6 of those individuals had to be excluded due to: 1) dysarthria and disphagia biasing scores on the LAST, so they didn’t really have aphasia even in the acute phase post-stroke (n=4); 2) extremely severe hearing loss (n=1); 3) no access to lesion data (n=1). This led to a final sample of 39 individuals with aphasia.

Healthy age-matched controls were recruited via flyers positioned in recreational community centers for elderly. The target age of healthy controls was gradually adapted based on the mean age of IWA included in the study.

